# Single-Cell Inference of Structural States Of Ribosomes

**DOI:** 10.64898/2026.08.29.747780

**Authors:** Euan Joly-Smith, Michael VanInsberghe, Kseniia Sarieva, Eugenio Marinelli, Robert M. van Es, Paula Sobrevals Alcaraz, Harmjan R. Vos, Amanda Andersson-Rolf, Hans Clevers, Alexander van Oudenaarden

## Abstract

Protein synthesis is dynamically regulated to control cell growth, differentiation, and stress responses. Recent single-cell sequencing methods can map ribosome positions on individual transcripts^1–4^, but cannot capture the global translational states that coordinate protein synthesis across the transcriptome. In contrast, methods that measure the global translational landscape, such as polysome profiling and cryogenic electron tomography^5^, lack either single-cell resolution or throughput. Here we introduce SCISSOR (Single-Cell Inference of Structural States of Ribosomes), a strategy that infers global translation activity in individual cells from the differential protection of ribosomal RNA (rRNA) against nuclease digestion. By integrating these protection signatures with the structure of the ribosome, SCISSOR resolves multiple ribosomal states and quantifies their abundance across thousands of individual cells. Applying SCISSOR reveals systematic variation in global translation across the cell cycle in human cells, as well as during the differentiation of murine intestinal stem cells into distinct epithelial lineages. These findings uncover principles of global translational regulation that are invisible to transcriptomic or ribosome-profiling assays, establishing a framework for studying global translation control at single-cell resolution.

---

Ribosome profiling relies on controlled nucleolytic digestion of cellular mRNAs to generate ribosome-protected fragments (RPFs)^6,7^. However, this digestion also affects other RNA species, most notably rRNA, which comprises a substantial fraction of sequencing reads in ribosome-profiling data sets^1,6^. These rRNA-derived reads are typically discarded as unwanted byproducts^6^. Recent computational studies, however, suggest that rRNA digestion patterns might be useful to explore ribosome protein stoichiometry^8^ and ribosome collisions^9^. Furthermore, a recent study in plants demonstrated that most rRNA fragments detected in bulk ribosome profiling originate near the solvent-exposed surface of the 80S ribosome^10^. We hypothesize that the rRNA fragments generated during a ribosome-profiling experiment are a rich source of information that allows a detailed inference of structural states of the ribosome. To disentangle cell-to-cell variation in these structural states from variation within individual cells, it is necessary to obtain rRNA digestion profiles at single-cell resolution. Here, we establish an integrated experimental and computational framework to enable Single-Cell Inference of Structural States Of Ribosomes (SCISSOR).

## rRNA digestion patterns in single cells accurately predict cell cycle state and intestinal cell type

In a ribosome-profiling experiment, the nuclease accesses rRNA regions differently depending on the structural state of the ribosome. For example, free subunits (40S and 60S) expose their intersubunit interfaces to digestion, while the 80S structure protects these interfaces (Fig. 1a). To experimentally quantify accessibility of rRNA to digestion at single-nucleotide resolution, we measure the cutting probability *P_cut_*defined as the ratio of the number of 5’ cut sites to the total coverage at each rRNA position in an individual cell (Fig. 1b). Positions with high *P_cut_* values indicate structurally exposed sites, while low *P_cut_* values correspond to protected rRNA sites. Crucially, in a single cell containing a mixture of ribosomal states, *P_cut_* will be determined by the relative abundance of these states and the ability of each state to protect the rRNA from digestion. To assess whether *P_cut_* varies between individual cell states, we first generated scRibo-seq libraries from human RPE-1 hTERT FUCCI cells^11^ collected from distinct states of the cell cycle and digested with a range of MNase concentrations spanning more than four orders of magnitude (*Methods*). We collected interphase (*n = 2152*) cells, contact-inhibited G0 cells (*n = 1166*) and cells in mitosis (*n = 360*). We first performed dimensionality reduction using uniform manifold approximation and projection (UMAP) by constructing a count matrix of reads aligning to protein-coding transcript sequences. As expected, a clear separation between the different cell cycle stages is observed (Extended Data Fig. 1a-d), as was previously reported^1^. Strikingly, when constructing a UMAP using *P_cut_* values derived solely from rRNA, cells segregated into distinct regions corresponding to cell cycle states. This indicates that rRNA protection against digestion is not constant during the cell cycle and that individual cell cycle states have distinct *P_cut_* profiles (Fig. 1c). To strengthen the generality of this observation, we generated scRibo-seq data from murine intestinal epithelial cells across two independent batches and performed analogous analyses (Extended Data Fig. 1e-h). Importantly, populations of enterocytes, progenitor cells, goblet cells, and enteroendocrine cells segregated in UMAP space based solely on their *P_cut_* profiles, demonstrating that rRNA digestion patterns alone are sufficient to distinguish major intestinal cell types (Fig. 1d).

**Figure 1.**
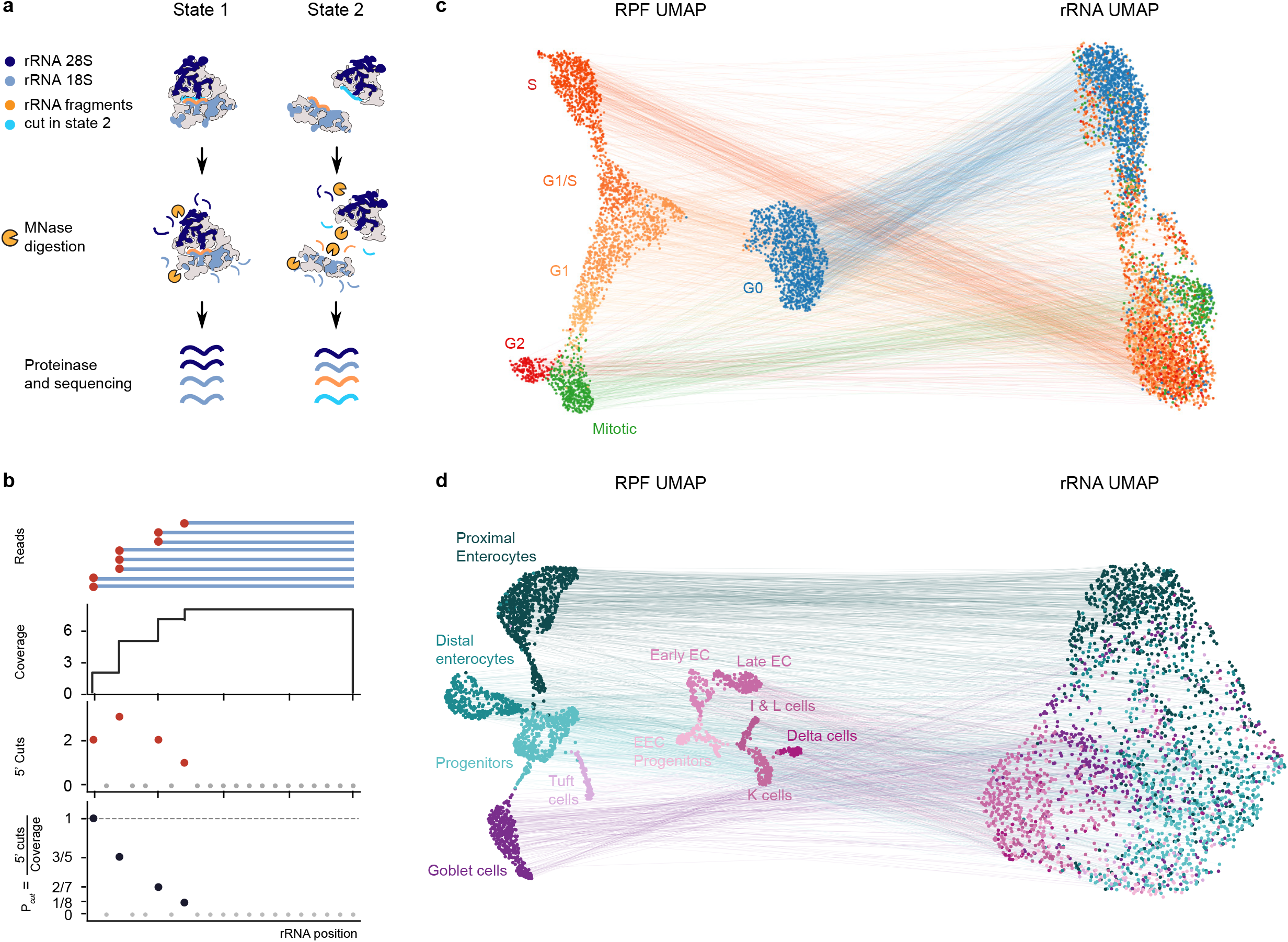
Single-cell rRNA cutting profiles encode cell-cycle and cell-type identity. **a,** Schematic illustration of conformation-dependent MNase digestion of ribosomal structures. MNase accessibility varies between different ribosomal conformations (illustrated here as assembled 80S ribosomes in State 1 versus open 40S and 60S subunits in State 2), resulting in differential digestion patterns. **b,** Definition and calculation of the *P_cut_* metric for quantifying rRNA protection. *P_cut_* at position *x* along the rRNA is defined as the ratio of read starts (cuts) to total coverage at that position. **c,** UMAPs generated from ribosome-protected fragment (RPF) counts and rRNA cutting profiles in RPE1-FUCCI cells. Cells from multiple MNase digestion levels were integrated before UMAP projection. Each point represents an individual cell, colored by cell-cycle phase (G0, G1, G2, mitotic, S). Left: UMAP based on normalized RPF counts per gene. Right: UMAP based on *P_cut_* values across rRNA positions. Both representations separate cells according to cell-cycle state. **d,** UMAPs generated from ribosome-protected fragment (RPF) counts and rRNA cutting profiles in mouse intestinal epithelium. Data from two independent batches were integrated before UMAP projection. Each point represents an individual cell, colored by cell type. Left, UMAP based on normalized RPF counts per gene. Right, UMAP based on *P_cut_* values across rRNA positions. Both representations separate major intestinal cell types. Together, these results demonstrate that single-cell rRNA cutting profiles contain sufficient information to distinguish biological cell states and cell types.

### Perturbation of ribosomal states reveals the digestion signatures of individual ribosome subunits

Having demonstrated that rRNA digestion profiles differentiate cell cycle states and intestinal cell types, we next aimed to find positions that are differentially accessible to digestion between fully assembled ribosomes and individual ribosomal subunits. To this end, we perturbed ribosomal states using translation inhibitors with well-documented effects on the ribosome. We exposed RPE-1 hTERT FUCCI cells to either 0.1 mg ml^-1^ puromycin (PURO) or 20 μM homoharringtonine (HARR) for 15 minutes. PURO is a translation inhibitor that mimics aminoacylated Tyr-tRNA and incorporates into the growing protein chain^12^. However, it cannot participate in the subsequent transesterification reaction, leading to premature termination of translation and ribosomal subunit release from the mRNA^13^. HARR, in turn, blocks early elongation following translation initiation^7,14^ while elongating ribosomes continue to completion, run off transcripts, and dissociate into subunits. As a result, transient treatment with either PURO or HARR increases the proportion of free ribosomal subunits in the cytoplasm. Consistent with the mechanism of action of both drugs, RPF coverage across coding sequences peaked near the start codon and declined sharply thereafter (Extended Data Fig. 2a). An RPF-based UMAP separated cells by treatment condition while also capturing their distribution across the unperturbed cell cycle (Extended Data Fig. 2b-d). In contrast, whereas HARR-and PURO-treated cells displayed distinct clusters in the RPF-based UMAP, an rRNA *P_cut_*-based UMAP revealed that PURO-and HARR-treated cells cluster together, clearly distinct from untreated cells, indicating similar rRNA digestion profiles under both treatments. (Fig. 2a). To characterize changes in rRNA digestion profiles, we performed differential digestion analysis at each rRNA position between conditions, analogous to differential gene expression analysis. We identified 399 and 318 differentially digested positions (DDPs) for HARR versus CTRL and PURO versus CTRL comparisons across concatenated 45S pre-rRNA and 5S rRNA, respectively. Figure 2b displays representative examples of DDPs. DDPs exhibiting increased digestion upon drug treatment significantly overlapped (Fisher’s exact test, p-value < 0.05) between the HARR and PURO treated cells, with 57.4% (28S) and 45.1% DDPs (18S) shared between comparisons (Fig. 2c, Extended Data Fig. 3e). Mapping DDPs onto a 3D cryo-EM structure of the ribosome (PDB: 6QZP) revealed enrichment at the intersubunit interface (Fig. 2d).

**Figure 2.**
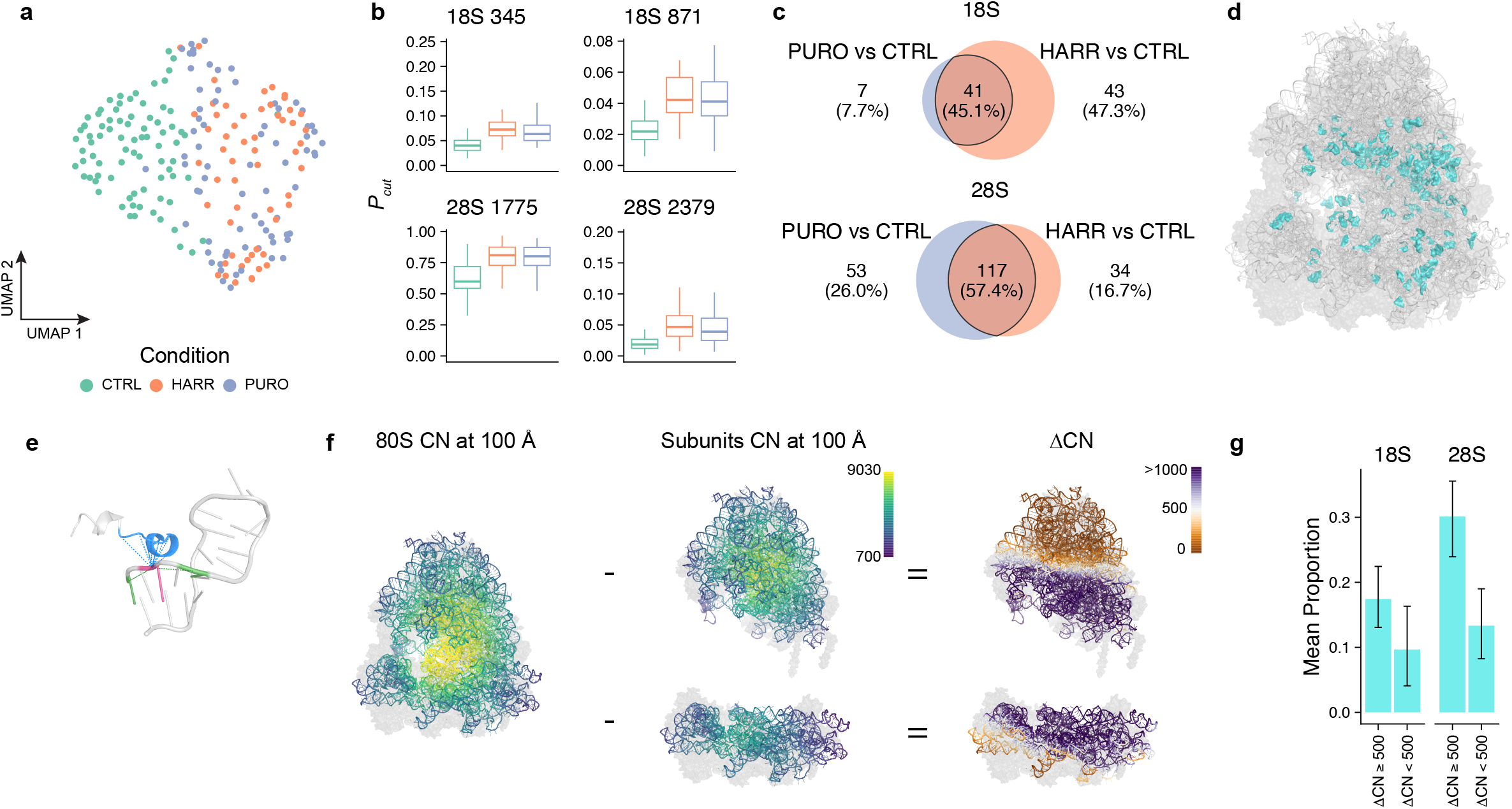
Translation inhibitors modify ribosomal state distribution. **a,** UMAP (n = 209 cells) of RPE1-FUCCI cells based on 5’ probability of cut. RPE-1 FUCCI cells were treated with either puromycin (PURO) at 0.1 mg/ml or homoharringtonine (HARR) at 20 µM for 15 minutes. Control (CTRL) cells were exposed to DMSO for the same duration. **b,** Differentially digested rRNA positions with the largest positive effect size difference shared between comparisons (PURO vs CTRL, HARR vs CTRL; significance defined as: absolute logit difference > 0.3, adjusted p-value < 0.05) between both comparisons). **c,** Venn diagrams showing the intersection of differentially digested positions between two comparisons (PURO vs CTRL, HARR vs CTRL) for 18S and 28S separately. Only positions with a positive logit difference are shown (logit difference > 0.3, adjusted p-value < 0.05). **d,** 3D representation of differentially digested positions (highlighted in teal on a cryo-EM model of the human ribosome) shared between comparisons from Fig. 2c. **e,** Schematic illustrating the concept of the contact number. For illustration purposes, nucleotides 189-209 of 18S and amino acids 137-147 of RPS8 were isolated. Contact number (CN) was defined as the number of neighboring nucleotides or amino acid for which at least one heavy atom lies within 12 Å of nucleotide 191’s P atom (pink). Immediately adjacent nucleotides (i ± 1 for illustration purposes, i ± 3 for the actual quantification) were excluded, so the CN reflects only tertiary RNA and protein contacts. **f,** Schematic illustrating defining the intersubunit interface through CN. First, the CN at 100 Å was calculated for the 80S ribosome (left) and individual subunits (60S and 40S, middle) separately. ΔCN (right) was calculated as the per-nucleotide difference in CN value between 80S and the respective individual subunit position. A ΔCN value of 500 or greater was used as a cutoff to define the intersubunit interface. **g,** Proportion of differentially digested positions with positive effect size among positions belonging to the intersubunit interface (ΔCN ≥ 500) and outside of the intersubunit interface (ΔCN < 500). Mean and error bars were obtained by bootstrapping differential digestion analysis by resampling cells 100 times. Error bars represent a bootstrap 95% confidence interval (1.96 ⤫ SD).

To systematically assess this enrichment, we sought an unbiased approach to assign nucleotides to the intersubunit interface. We therefore employed the concept of contact number (CN), defined in protein structural biology as the number of Cα atoms within a given radius of the target Cα atom, excluding immediate neighbors^15,16^. Conceptually, this metric characterizes how densely a certain amino acid is surrounded by other amino acids within the protein tertiary structure. We modified this metric to quantify CN for P-atoms of nucleotides by counting the number of surrounding nucleotides and amino acids within a defined radius (Fig. 2e). With the ribosome being 250-300 Å in diameter, the CN at 100 Å radius provides an impression of the gross ribosome architecture (Fig. 2f). To identify the intersubunit interface, we computed CN for individual subunits by removing the opposing subunit from the 80S structure. ΔCN, defined as the difference between 80S and subunit CN, highlights the intersubunit interface (Fig. 2f). We next quantified the proportion of DDPs among positions with ΔCN ≥ 500 (intersubunit interface) and ΔCN < 500 (outside of the intersubunit interface). The proportion of DDPs was approximately three times higher for 28S within the intersubunit interface than outside with the same tendency for 18S, indicating enrichment of DDPs at the intersubunit interface (Fig. 2g). Collectively, these results demonstrate that rRNA digestion profiles from single cells carry information about ribosomal states, and that perturbations of ribosomal states reveal DDPs distinguishing free-floating ribosomal subunits from fully assembled ribosomes.

### Deconvolving ribosomal states from naturally occurring cell-to-cell variability

Next, we asked whether similar structural information could be extracted in unperturbed populations by leveraging naturally occurring variability between single cells. In any given cell, ribosomes exist as a mixture of structural states, each with distinct MNase cutting profiles and abundances. The cuts observed at each rRNA position reflect both the relative abundance of ribosomal states and their position-specific cleavage affinities. To capture this quantitatively, we developed an analytical framework that models MNase cleavage while accounting for both ribosomal state heterogeneity and technical noise. In this framework, the observed cuts at each rRNA position arise from contributions of multiple ribosomal states with distinct cutting probabilities and abundances. Additional factors, such as cell-to-cell variation in sequencing efficiency, sequence-specific biases, and size selection due to poorly mapped short reads, also influence the observed counts (*Methods*). A schematic of the overall analysis pipeline is shown in Fig. 3a. From this general model, we analytically show (*Methods*) that whenever two positions *x* and *y* in the rRNA have the same relative cutting probabilities across all ribosomal states, then their normalized covariance with a third position *z* is equal to: *η_xz_ = η_yz_*, where *η_xy_* is the normalized covariance between the relative frequency of cuts at position *x* and *y* computed over all individual cells (Fig. 3a). Cuts at positions with similar accessibility across ribosomal states co-vary in a similar way across cells. High *η_xy_* values indicate positions that are likely cut in the same ribosomal states, providing a means to group rRNA positions. Building on this insight, we clustered rRNA positions based on their *η_xy_* values to identify groups of co-varying sites (Fig. 3b, *Methods,* Supplementary Note). The resulting heatmap reveals clusters with structural features: some clusters correspond to regions on the ribosomal surface with low contact number, while others are enriched in pre-rRNA regions. These features are quantified in Fig. 3c, which shows the distribution of contact numbers across positions in each cluster (Fig. 3c, violin plots) and the fraction of positions within pre-rRNA regions (Fig. 3c, histogram).

**Figure 3.**
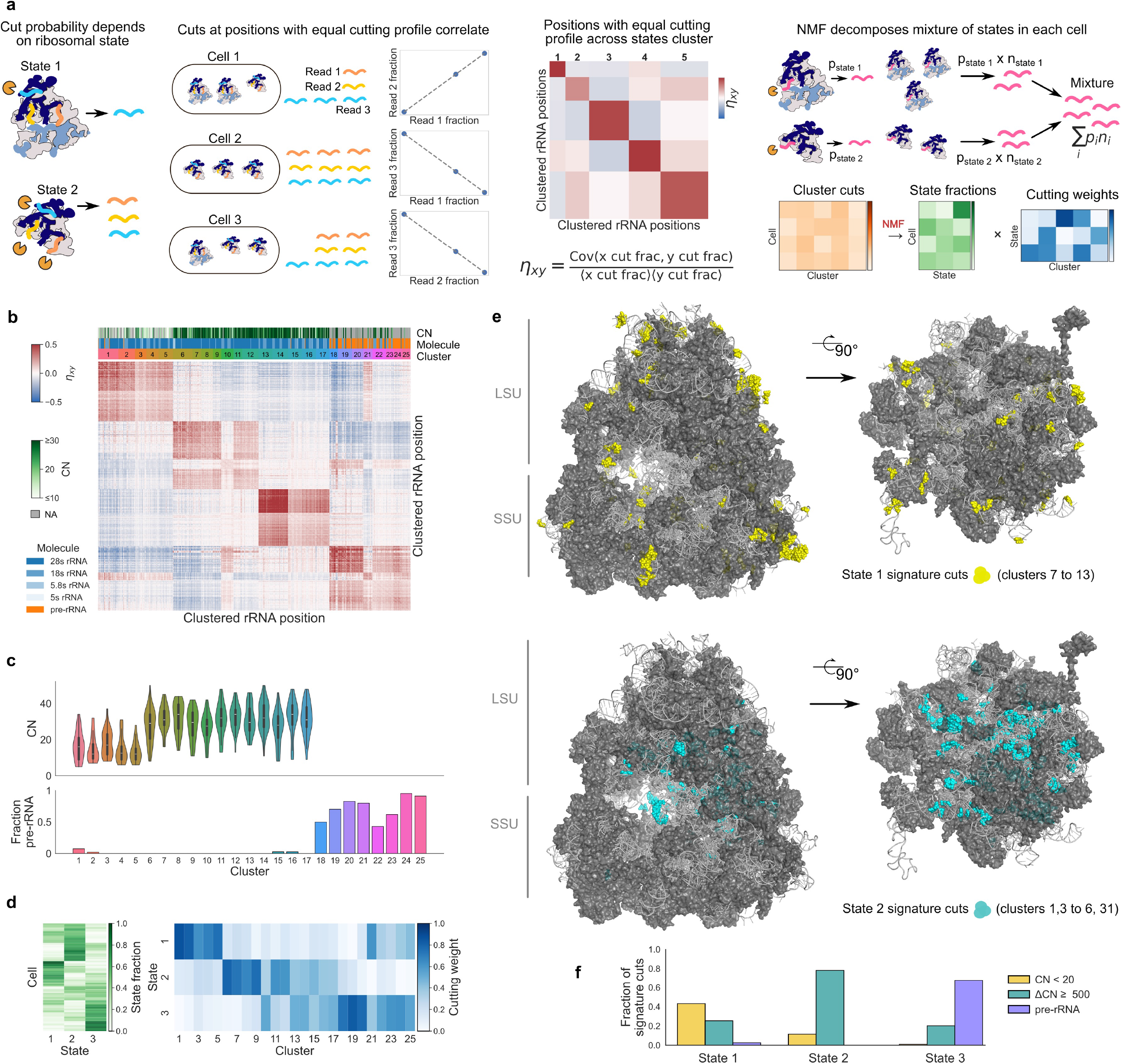
Deconvolution of ribosomal states from single-cell rRNA cutting profiles. **a,** Schematic overview of the analysis pipeline: ribosomes in each cell exist as a mixture of states with distinct MNase cutting profiles; positions with similar state-specific cutting profiles co-vary across cells; clustering groups co-varying positions; NMF extracts state-specific cutting signatures and relative per-cell state abundances. **b,** Heat map of measured *η_xy_* values illustrating clustered rRNA positions. Three color bars show the CN with radius 12 Å for each position, the molecule identity (pre-rRNA, 18S, 28S, or 5.8S rRNA), and the cluster assignment. Positions on the ribosomal outer surface with low CN cluster together, as do positions in pre-rRNA. **c,** Violin plots of CN across positions in each cluster, and a histogram showing the fraction of positions in pre-rRNA regions, highlighting structural features of clusters. **d,** NMF output matrices: per-cell state fractions (left) and cutting weights (right). The cutting weights indicate the fraction of reads in each cluster derived from each state. Clusters in which a state contributes ≥2-fold more than any other are defined as *signature cuts* for that state. **e,** Cryo-EM models of the human ribosome with signature cuts highlighted. Top: state 1, showing surface-enriched signature cuts. Bottom: state 2, showing interface-enriched signature cuts. **f,** Fraction of signature cuts for each state with CN < 20 (with 12 Å radius), ΔCN ≥ 500 (with 100 Å radius), or mapping to pre-rRNA. State 1 consistently exhibits enrichment for cuts with CN < 20, corresponding to regions near the ribosomal outer surface. State 2 exhibits enrichment for cuts with ΔCN ≥ 500, corresponding to regions near the intersubunit interface. State 3 consistently exhibits enrichment in pre-rRNA. The above analysis was done on cells at MNase digestion level 0.39 U. These results are reproducible across increased MNase digestion levels (Extended Data Fig. 3).

Positions within the same cluster share similar cutting probabilities across ribosomal states, therefore each position reflects approximately the same linear combination of underlying state abundances. Summing cuts across all positions in a cluster for each cell amplifies this signal while reducing noise, giving a cell-by-cluster matrix in which each entry represents the total cuts for a given cluster in a given cell. To decompose these unknown linear combinations, we applied non-negative matrix factorization (NMF) to this matrix, inferring both the characteristic cutting profiles of each ribosomal state (cutting weights matrix, blue) and their relative abundances across cells (state fraction matrix, green; Fig. 3d). Note, the cutting weights matrix theoretically represents the probability of a cut in cluster *i* given ribosomal state *j*, but its scale is non-unique. We therefore rescaled the cutting weights by the average contribution of each state across clusters, which results in the scale-independent fraction of reads in cluster *i* derived from state *j*. For a given ribosomal state, clusters in which this fraction was at least 2-fold higher than for all other states were defined as *signature cuts* for this state of interest. This provides a basis for interpreting the structural identity of each ribosomal state.

Using this framework, we consistently recovered three reproducible ribosomal states across different MNase concentrations (Extended Data Fig. 3a-c). One displayed surface-enriched cuts, consistent with intact 80S ribosomes (signature cuts highlighted on a cryo-EM structure in Fig. 3e, top). A second exhibited interface-enriched cuts, corresponding to free subunits (Fig. 3e, bottom). A third showed strong signals within pre-rRNA regions, indicative of biogenesis intermediates (Fig. 3f). At very low MNase concentrations, however, the resulting states were not readily interpretable (Extended Data Fig. 4a-e). These results demonstrate that even in unperturbed populations, rRNA cutting profiles encode rich structural information, enabling quantitative mapping of ribosomal states in single cells. This strategy can be readily applicable to heterogeneous cellular systems such as cell cycle states or complex tissues, such as mouse intestine.

### Ribosomal states in single human cells change during the cell cycle

Having developed two independent approaches to decipher relative proportions of ribosomal states between different cellular states and types, we next aimed to compare their predictions of the dynamics of ribosomal states during the human cell cycle. We use the following two metrics. The first metric (Fig. 4a) was derived from the summed 5′ cut signal at DDPs shared between HARR and PURO treatments as explained in Fig. 2, divided by the total RPF count per cell. This metric thus reports on the ratio between free ribosomal subunits and translating ribosomes. The second metric originates from the NMF approach (Fig. 4b) as explained in Fig. 3. Both approaches yielded a qualitatively similar result that G0 and mitotic cells have a higher proportion of free-floating subunits than interphase cells (Fig. 4a-b). This is consistent with the existing literature for both cell states. Indeed, G0 cells downregulate translation^17^ with concurrent decrease in polysome abundance^18,19^. Similarly, mitotic cells downregulate translation by about 35%^20^. However, published studies reporting polysome profiles in mitotic cells have produced inconsistent results. Some studies report that the reduction in global translation rates in mitotic cells is a result of ribosomes dwelling on transcripts, yielding polysome profiles that are similar to interphase cells^21–23^. Others report the reduction in the proportion of polysomes^24–27^ with a simultaneous increase in signal from individual subunits^24^. Hence, to validate our observations, we performed Mega-SEC^28,29^, a technique that allows resolving ribosomal complexes using size-exclusion chromatography (Fig. 4c). We used protein mass spectrometry (MS) to validate fraction composition. MS revealed 9,265 individual proteins across samples with overall stable total iBAQ per sample (Extended Data Fig. 5a-b). Fractionation was driving principle component analysis (Extended Data Fig. 5c) with fractions 8 and 9 being extraribosomal (Extended Data Fig. 5d). The relative iBAQ (riBAQ) ratio of large (LSU) to small (SSU) ribosomal subunit proteins enabled assignment of fractions to polysomal, 80S, mixed 60S/80S, and free 40S populations (Fig. 4d). We next examined changes in ribosomal state abundance across cell-cycle stages. G0 and mitotic cells exhibited reduced polysomal peak areas relative to interphase cells (Fig. 4e, Extended Data Fig. 5e,f). As no fraction corresponding to a pure 60S population was identified, the proportion of 60S ribosomes was estimated from fitted polysome, monosome, and 60S peaks using absorbance measurements together with proteomic estimates of subunit composition and a Beer–Lambert-based mixture model (Fig. 4d, Extended Data Fig. 5e,f, Supplementary Note). We found a higher proportion of 60S in G0 and mitotic than in interphase cells, hence confirming the higher proportion of individual subunits in these cells, as predicted by SCISSOR (Fig. 4f).

**Figure 4.**
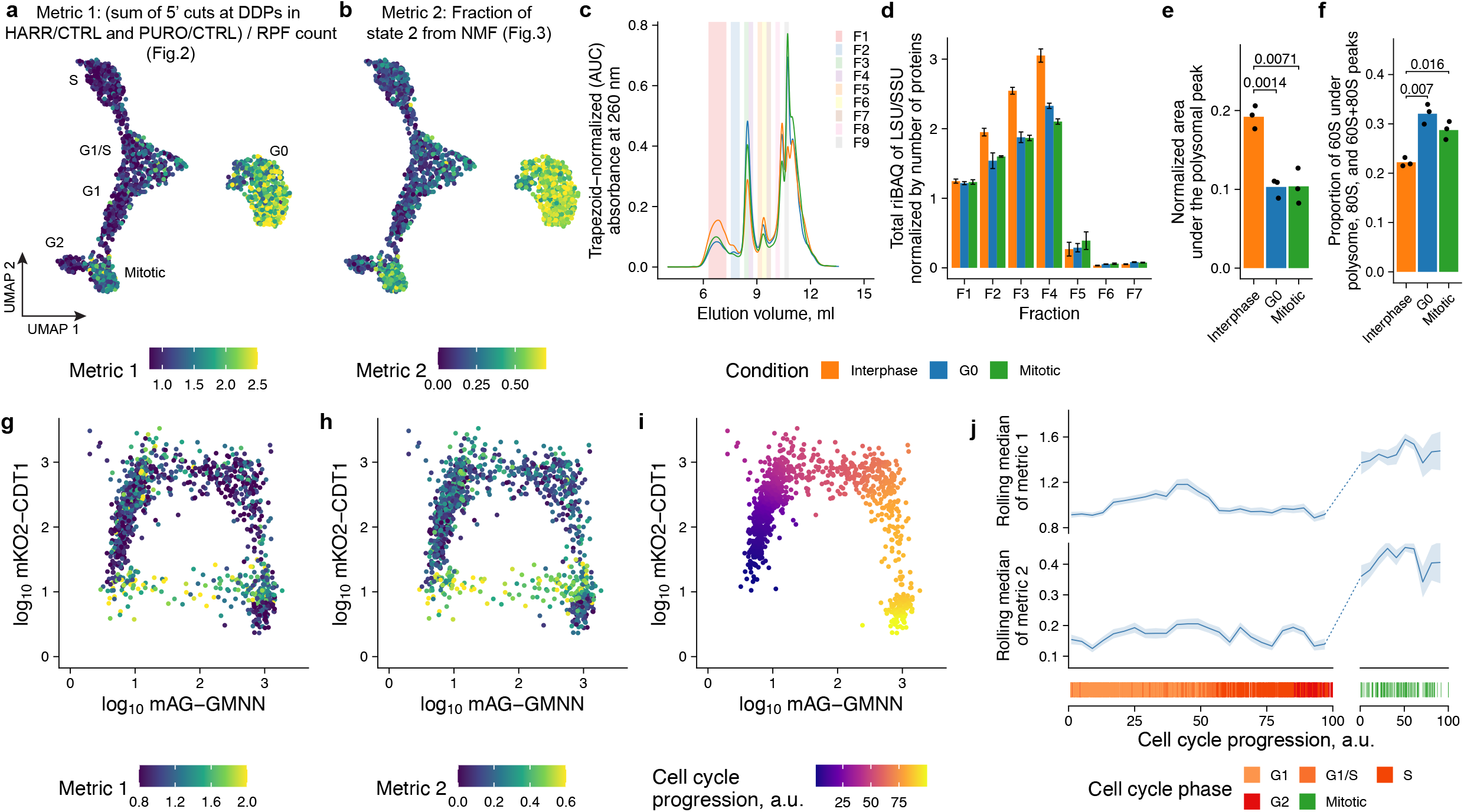
Ribosomal state distribution is dynamic across the cell cycle. **a,b,** UMAP of RPE1-FUCCI cells based on RPFs. Cells treated with multiple MNase concentrations are shown. **a,** Color represents the sum of 5’ cuts at the shared DDPs between HARR/CTRL and PURO/CTRL comparisons from figure 2, divided by the number of RPF within CDS per cell. Only positions with the mean number of cuts > 10 in interphase cells were used. To correct for differences in proportions across MNase concentrations, all values were normalized to the average in interphase cells for a given MNase concentration. **b,** Color represents the fraction of state 2 as assessed by NMF. **c,** Representative Mega-SEC polysome profiling absorbance (260 nm) traces from RPE1-FUCCI interphase, G0 and mitotic cells. Fractions collected for protein mass spectrometry are highlighted. **d,** Relative iBAQ (riBAQ) ratios of large (LSU) to small (SSU) ribosomal subunit proteins across fractions. Fraction 1 exhibited an LSU/SSU ratio of approximately 1, consistent with polysomes, whereas fractions 5–7 were enriched for SSU proteins, indicating free 40S subunits. Fractions 2–4 displayed intermediate LSU/SSU ratios. Fractions 3 and 4 were thus assigned to mixed 60S/80S populations and fraction 2 to 80S ribosomes. As ribosomal proteins contributed only a small fraction of total protein abundance in fractions 5–7 (Extended Data Fig. 5d), these fractions were excluded from subsequent subunit-composition analyses. **e,** Comparisons of area under the curve for polysomal peak from c. Curves were obtained by fitting exponentially-modified Gaussian distributions. **f,** Comparisons of proportion of 60S across polysomal, 80S, and 60S+80S fractions. **e,f,** Bars represent the mean, dots represent individual experiments (n = 3, two-sided pairwise t-test). **g,h,** Scatter plots of the FUCCI markers with colors denoting values for metric 1(**g**) and metric 2 (**h**). Only interphase and mitotic cells are shown. i, Scatter plots of the FUCCI markers with colors denoting time along cell cycle progression as calculated by ERA for interphase cells. j, left panels, rolling median plots of metric 1 (top) and metric 2 (bottom) over cell cycle stage progression in interphase cells from i. Right panels, rolling median plots of metric 1 (top) and metric 2 (bottom) over cell cycle progression in mitotic cells. The rolling median window was set to 8 and 20 arbitrary units (a.u.), and the step size to 2 and 10 a.u for interphase and mitotic cells, respectively. Lines represent the rolling medians, ribbons represent the standard error of the median absolute deviation (MAD-SE). Bottom strip plots show assignment of cells to cell cycle stage progression.

Next, we wanted to analyse the dynamics of the relative amount of free subunits along the cell cycle (Fig. 4g for metric 1 and Fig. 4h for metric 2). Using Ergodic Rate Analysis^30^ we could assign pseudotime to each cell along a cell cycle stage, separately for interphase and mitotic cells (Fig. 4i, Extended Data Fig. 5g). Using these pseudotime assignments, we found a moderate increase in both metrics in late G1 phase cells and a sharp increase as cells enter mitosis (Fig. 4j). In addition to changes in free subunit proportion across cell cycle, we noticed that the fraction of ribosome biogenesis state 3 derived from NMF analysis was lower in both G0 and mitotic cells with the same tendency observed for the intact 80S ribosome state 1 (Extended Data Fig. 5h,i). Collectively, we observed large changes in ribosome states during the human cell cycle, revealing an additional layer of global regulation of gene expression.

### SCISSOR reveals cell-type-specific ribosomal states in the intestinal epithelium

Having established that ribosomal state composition changes dynamically during the human cell cycle, we next asked whether SCISSOR could be applied to a complex mammalian tissue. We applied the analytical framework described in Fig. 3 to infer ribosomal states during the differentiation of murine intestinal stem cells into distinct epithelial lineages. Clustering of rRNA positions based on co-variation of cuts across cells, followed by non-negative matrix factorization, resulted in a decomposition into five ribosomal states (Extended Data Fig. 6a,b). Mapping signature cuts onto a cryo-EM structure of the mouse ribosome (PDB: 7CPU) and the secondary structure of the mouse 28S rRNA revealed distinct structural features for each state. State 1 displayed cuts enriched on the ribosomal surface and on expansion segments, consistent with intact 80S ribosomes (Fig. 5a). State 2 showed enrichment along the intersubunit interface and within the interior of the ribosome, consistent with free subunit state (Fig. 5b). State 3 showed strong enrichment in pre-rRNA regions, indicative of biogenesis intermediates (Fig. 5d). In addition, we identified a fourth state characterized by increased cuts in multiple expansion segments, including ES7L, ES27L, ES30L, ES31L and ES39L, exceeding those observed in state 1 (Fig. 5c). Lastly, a fifth state did not display a clear structural signature or cell-type-specific enrichment (Extended Data Fig. 6g,h). We therefore did not assign a biological interpretation to this state and excluded it from downstream analyses of state fractions. These states were reproducible in an independent experimental batch (Extended Data Fig. 6a-f).

**Figure 5.**
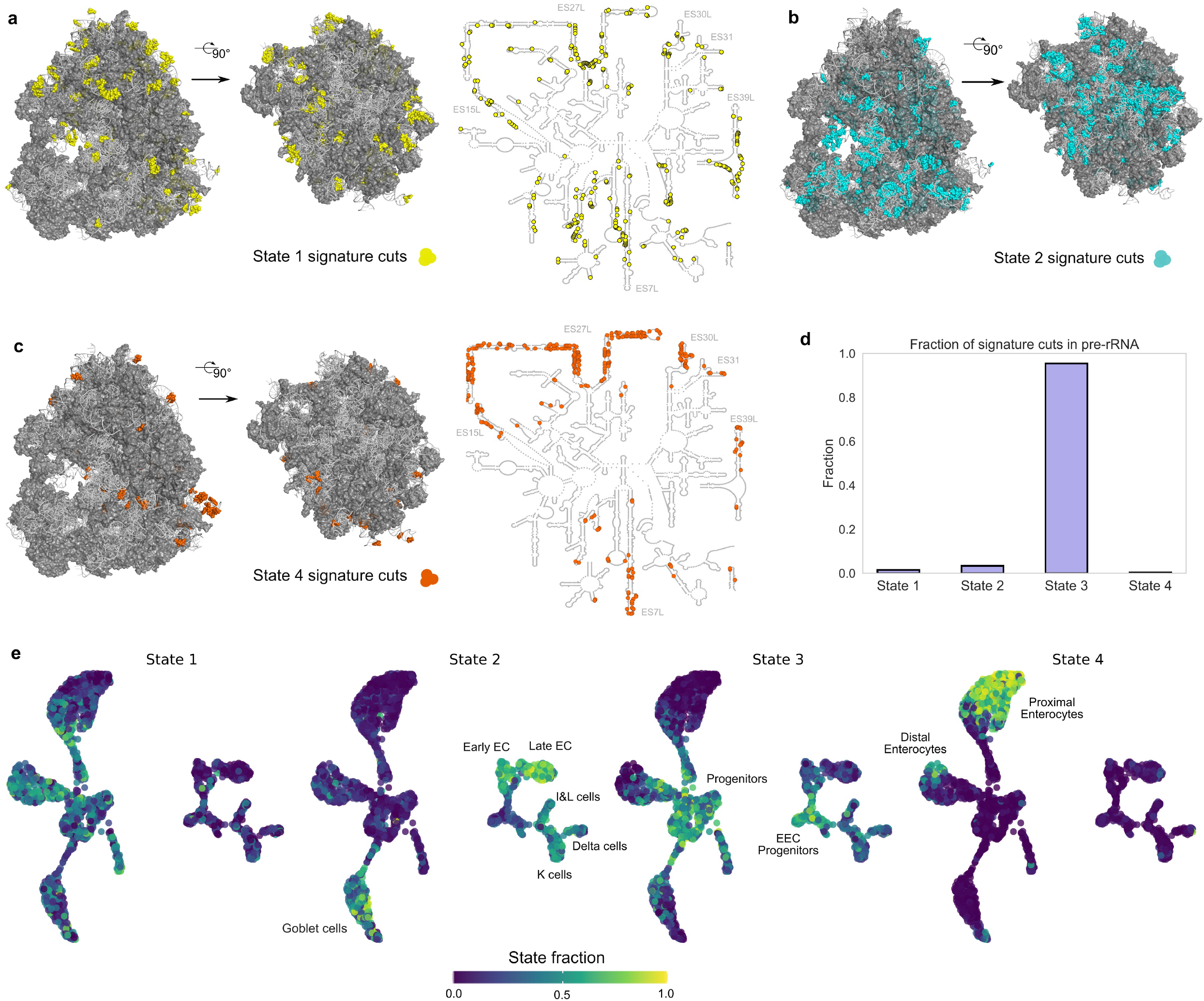
Structural ribosomal states and cell-type-specific global translation in the murine intestine. **a,** Signature cuts for state 1 from one representative experimental batched mapped onto the mouse ribosome and the mouse 28S rRNA secondary structure. The intact 80S ribosome is shown (left), alongside a view of the isolated 60S subunit with the 40S subunit removed and reoriented to expose the intersubunit interface (right). Cuts are enriched on the outer surface, with additional contributions in expansion segments in the secondary structure representation. This pattern is consistent with intact 80S ribosomes. **b,** Signature cuts for state 2. Cuts are enriched within the ribosomal interior and along the intersubunit interface, consistent with free subunits. **c,** Signature cuts for state 4. Cuts are enriched in expansion segments (ES7L, ES27L, ES30L, ES31L and ES39L), defining an expansion segment–associated ribosomal state. **d,** Fraction of signature cuts of each state that are localized in pre-rRNA. The majority of state 3 signature cuts are localized in pre-rRNA regions, indicating that this state consists of ribosome biogenesis intermediates. **e,** UMAP embedding of intestinal cells derived from RPFs (Fig. 1d), overlaid with inferred ribosomal NMF state fractions (left to right: states 1–4) from two independent batches (see *Methods* for batch integration procedure). Distinct epithelial populations exhibit characteristic ribosomal state compositions. The free subunit state (state 2) is enriched in secretory cells. The biogenesis intermediate state (state 4) is elevated in progenitor cells. The expansion segment–associated state (state 3) is enriched in proximal and distal enterocytes.

Next, we examined how the four identified ribosomal states are distributed across intestinal cell types. We projected inferred state fractions onto the UMAP derived from ribosome-protected fragments (RPFs) (Fig. 1d; Fig. 5e). Distinct epithelial populations exhibited characteristic ribosomal state compositions. In particular, state 2, the free subunit state, was enriched in secretory cells, whereas the biogenesis intermediate state was elevated in progenitor cells, consistent with increased ribosome production in dividing cells. In contrast, the expansion segment–associated state was preferentially enriched in both proximal and distal enterocytes.

These results extend our observations from the cell cycle in human tissue culture cells to a complex heterogeneous tissue. SCISSOR enables direct quantification of global translation states in single cells and reveals their coordinated variation across epithelial lineages.

## Discussion

By leveraging rRNA digestion patterns that are typically discarded in ribosome-profiling experiments, SCISSOR provides a general framework to infer global translational states directly in single cells. Unlike transcriptomic approaches, which primarily capture the potential for protein production, or conventional ribosome profiling, which reports ribosome occupancy on individual transcripts, SCISSOR resolves coordinated shifts in the overall structural and functional landscape of the translational machinery itself. Our analyses reveal that these global ribosomal states are tightly coupled to major cellular transitions, including cell-cycle progression and epithelial differentiation in the intestine, and uncover distinct structural signatures associated with free subunits, assembled ribosomes, ribosome biogenesis intermediates, and expansion-segment-associated conformations. Further work is required to decipher the molecular drivers of these different conformations.

More broadly, the ability to quantitatively map ribosomal states across thousands of individual cells opens opportunities to study translational regulation in development, regeneration, stress adaptation, aging, and disease. Because SCISSOR builds directly on standard ribosome-profiling workflows, it can in principle be integrated with existing and future single-cell translatomic technologies, providing a scalable route toward a systems-level understanding of global translation control in heterogeneous tissues.

## Materials and Methods

### Cell culture and dissociation

RPE-1 hTERT FUCCI cells were a kind gift of the Medema lab (The Netherlands Cancer Institute, Amsterdam, The Netherlands). Cells were cultured in DMEM/F12 supplemented with 10% FBS (Gibco), 1× GlutaMAX (Gibco), and 1× Pen-Strep (Gibco) at 37 °C with 5% CO2. For the translation inhibitor treatment experiments, 4.45 × 10^5^ cells were plated in a 10-cm plate. 36 h later, cells were washed with PBS0, and cells were treated with puromycin at a final concentration of 0.1 mg ml^-1^, homoharringtonine at 20 μM, or dimethyl sulfoxide (DMSO) at the equivalent dilution (1:1000) as a control diluted in cell culture medium. After 15 min, the cells were collected by trypsinization (TrypLE, Gibco). Cells were washed once and resuspended in the cold culture medium containing their respective treatment compound and DAPI as a viability stain. Cells were passed through a 20-μm mesh before sorting.

For the cell-cycle stages experiments, cells were treated as described before^1^. Briefly, for interphase cells, 4.45 × 10^5^ cells were seeded in a 10-cm plate and collected 36 h later. For mitotic cells, 1 × 10^6^ cells were seeded in a 15-cm plate and collected 36 h later. For G0 cells, 3 × 10^6^ cells were seeded in a 10-cm plate and collected 72 h later. Experiments were planned so that all experimental conditions could be collected simultaneously. For further cell sorting, cells were washed once with PBS0 and collected by trypsinization (for interphase and G0 conditions) or mitotic shake-off and spun down at 600 g for 3 min. Cells were further resuspended in complete cold media supplemented with DAPI and passed through a 20-μm mesh before sorting.

For Mega-SEC, Mitotic cells were collected by first changing the media to a fresh one to remove dead cells and debris, followed by mitotic shake-off, and spinning down at 600 g for 3 min. Cells were resuspended in cold PBS0, spun down at 1,000 g for 5 min at 4 °C, and the pellet was snap-frozen in liquid nitrogen. G0 and Interphase cells were washed twice with cold PBS0 on ice, collected by scraping in cold PBS0, and spun down at 1000 g for 5 min at 4 °C. The pellet was snap-frozen in liquid nitrogen, and samples were further processed as described. All samples were further thawed on ice, and Mega-SEC lysis buffer was added (20 mM HEPES-NaOH pH 7.5, 130 mM NaCl, 10 mM MgCl_2_, 5 % glycerol, 1 % CHAPS, 2.5 mM dithiothreitol (DTT), 2 U μl^-1^ RNaseIN Plus [Promega], 1× cOmplete EDTA-free Protease inhibitor [Roche], 0.11 mg ml^-1^ cycloheximide [Sigma-Aldrich]). Samples were incubated for 15 min at 4 °C on a rotating wheel, centrifuged at 16,049 g for 20 minutes at 4 °C, and the supernatant was passed through a 0.45 μm Ultrafree-MC HV centrifugal filter units (Merck Millipore) by centrifugation at 12,000 g for 5 minutes at 4 °C to remove remaining lipids and impurities. RNA concentration was measured by Qubit, and samples were diluted to the same RNA concentration with the lysis buffer.

Mouse intestinal epithelial cells were isolated from Neurog3Chrono mice, closely following the methods outlined by Gehart et al^31^. In brief, mouse small intestines were collected, cleaned, flushed with PBS0, and separated into proximal and distal sections. Tissue pieces from each region were opened and gently scraped with a glass coverslip to collect the villi fractions. The remaining tissue pieces were then washed in cold PBS0 before transferring to PBS0 with 2 mM EDTA (Gibco), incubated at 4 °C for 30 min on a roller, and then vigorously shaken to release crypts. Villi and detached crypts were pelleted, resuspended in warm TrypLE Select (Gibco), incubated at 37 °C for 2 minutes, and mechanically disrupted by pipetting to generate single-cell suspensions. Single-cell suspensions were washed twice in Advanced DMEM/F12 (Gibco), strained with a 20-μm mesh, and resuspended in FACS media [Advanced DMEM/F12 (Gibco), 1 % FBS, 1× GlutaMAX (Gibco), 10 mM HEPES (Gibco)] and incubated for 10 min at RT. Cells were then stained in FACS media with anti-EPCAM (ThermoFisher 15380750) and 1 μg ml^-1^ DAPI for 30 min on ice, were washed 2 times in FACS media, resuspended in FACS media with 1 μg ml^-1^ DAPI, and passed through a 20-μm mesh for sorting.

### Mice

All mouse experiments were conducted under the project license AVD8010020151 granted by the Dier Experiment Commissie/Animal Experimentation Committee (DEC) or Central Committee Animal Experimentation (CCD) of the Dutch government and approved by the Hubrecht Institute Animal Welfare Body (IvD). The Neurog3Chrono allele was maintained on a mixed *Mus musculus* C57BL/6 background. Animals used in the experiments were between 8 and 22 weeks of age. Both male and female mice were used for the experiments. Mice were housed in open housing with 14:10 h light:dark cycle at 24 °C and 45–70 % relative humidity with food and water ad libitum. The intestines from three individuals were pooled together during cell dissociation; randomization and blinding were not performed.

### FACS

Single cells were index sorted into 384-well plates using a BD FACS Influx with the following settings: sort objective single cells, a drop envelope of 1.0 drop, a phase mask of 10/16, a maximum of 16 extra coincidence bits, drop frequency of 38 kHz, a nozzle of 100 μm with 18 PSI and a flowrate of approximately 100 events per s, which results in a minimum sorting time of approximately 5 min per plate. Doublets, debris, and dead cells were excluded by using the forward and side scatter and the DAPI channels, respectively.

For the hTERT RPE-1 FUCCI cells, fluorescence in the monomeric Azami-Green (mAG) and monomeric Kusabira-Orange 2 (mKO2) channels was measured and later used for data analysis.

For the mouse intestinal epithelial cells, cells were further gated to select for EPCAM+. Enteroendocrine cells were further gated using the dTomato and mNeonGreen reporters from the Neurog3Chrono reporter^31^.

### Library preparation for scRibo-seq

Libraries were prepared as previously described with minor modifications^1^. Briefly, single cells are sorted into wells containing lysis buffer. They were subjected to nucleolytic digestion, followed by proteolytic treatment. Libraries were constructed by a one-pot small-RNA protocol consisting of end repair, 3’ DNA adapter ligation, reverse transcription primer annealing, 5’ RNA adapter ligation, reverse transcription, and indexing PCR.

In detail, single cells were sorted into 384-well hardshell plates (Bio-Rad) preloaded with 6 μl of light mineral oil (Sigma-Aldrich) and 50 nl of lysis buffer (22 mM Tris-HCl pH 7.5, 16.5 mM MgCl₂, 5.5 mM CaCl₂, 165 mM NaCl [Sigma-Aldrich], 1.1 % Triton X-100, 2.2 U μl^-1^ RNaseIN Plus, 0.11 mg ml^-1^ cycloheximide). Plates were centrifuged at 2,000 g for 2 min at 4 °C, kept on wet ice during sorting, and either processed immediately or stored at –80 °C until further use. Before processing, plates were thawed and centrifuged again at 2,000 g for 2 min at 4 °C.

For the cell-cycle stages experiment, nucleolytic digestion was initiated by adding 50 nl of MNase (New England Biolabs, NEB) at variable concentrations (25 U diluted down in a 15-point 2× dilution series, and 0 U MNase) to each well, followed by incubation at 37 °C for 30 min. For the translation inhibitor experiment, a MNase concentration 3.125 U per reaction was used. Digestion was terminated by adding 50 nl of stop solution (0.0186 U μl^-1^ Thermolabile Proteinase K [NEB], 62 mM EGTA, 16.5 mM EDTA [Ambion], 697.5 mM guanidinium thiocyanate [Sigma-Aldrich]). Plates were incubated at 37 °C for 30 min, then 55 °C for 10 min, and held at 4 °C.

End repair was performed by adding 50 nl of end-repair mix (4.1× T4 RNA Ligase Buffer [NEB], 16.4 mM MgCl₂, 4.1 mM UTP, 1.37 U/μl T4 Polynucleotide Kinase [NEB], 0.82 U/μl RNaseIN Plus) and incubating the plates at 37 °C for 1 h. Next, 3’ adapters were ligated by adding 264 nl of ligation mix (1× T4 RNA Ligase Buffer [NEB], 1 μM pre-adenylated 3ʹ adapter [Integrated DNA Technologies, IDT], 35.5% PEG-8000 [Sigma-Aldrich], 0.1% Tween-20, 1 U μl^-1^ RNaseIN Plus, 21.3 U μl^-1^ T4 RNA Ligase 2 Truncated KQ [NEB]) and incubating the plates at 4 °C for 18 h. After 3’ adapter ligation, the reverse transcription (RT) primer was annealed by adding 50 nl of RT primer mix (5.2 μM RT primer [IDT], 13.5 μM ATP [NEB], 1% Tween-20) and incubating the plates at 65 °C for 1 min, 37 °C for 2 min, 25 °C for 2 min, and holding at 4 °C. 5ʹ adapter ligation was performed by adding 156 nl of 5ʹ ligation mix (1× T4 RNA Ligase Buffer, 30.75% PEG-8000, 0.1% Tween-20, 0.5 μM 5ʹ adapter [denatured at 70 °C for 2 min and snap chilled on ice, IDT], 1.25 U μl^-1^ T4 RNA Ligase 1 [Ambion]) and incubating the plates at 37 °C for 2 h and holding at at 4 °C. Samples were reverse transcribed by adding 770 nl of RT mix (1.88× RT Buffer [Thermo Scientific], 1.25 mM dNTPs [Promega], 0.1875% Tween-20, 1.875 U μl^-1^ RNaseIN Plus, 9.375 U μl^-1^ Maxima H Minus Reverse Transcriptase [Thermo Scientific]) and incubating the plates at 50 °C for 1 h, then 85 °C for 5 min, and holding at at 4 °C. Finally, libraries were amplified by indexing PCR by adding 150 nl of 20 μM forward primers (IDT) containing cell barcode and UMI and 3.2 μl PCR mix (1.5× Q5 Hot Start Master Mix [NEB], 0.15% Tween-20, 0.94 μM reverse index primer [IDT]) and incubating at 98 °C for 30 s, followed by 10 cycles of 98 °C for 15 s, 65 °C for 30 s, and 72 °C for 30 s, and finally at 72 °C for 5 min and holding at 4 °C.

Final libraries were stored at –20 °C until pooling. All liquid handling was performed using either a Nanodrop II (Innovadyne Technologies) or Mosquito (TTP Labtech) platform. Plates were centrifuged at 2,000 g after each liquid transfer step.

### Plate pooling and purification

After library construction, the plates were pooled, and libraries were purified. The contents of each plate were first collected in VBLOK200 reservoirs (Click Bio) by centrifuging at 2,000g for 2 min. The aqueous phase (∼1.9 ml per plate) was separated from the light mineral oil by centrifugation, and concentrated to approximately 500 μl using n-butanol (Sigma-Aldrich). Product was then cleaned up using AMPure XP beads (Beckman Coulter) that had been diluted 6× in bead binding buffer (20% PEG-8000, 2.5 M sodium chloride); diluted beads were added to the sample at a 2.1:1 ratio, and the final product was resuspended in 20 μl LoTET buffer (3 mM Tris-HCl pH 8.0 [Ambion], 0.2 mM EDTA pH 8.0, 0.1% Tween-20). Half of each of the cleaned-up library pools was then run on a 7% polyacrylamide gel at 300 V for ∼3 h, and gel lanes above ∼160 nt, corresponding to an insert size of more than 15 nt were excised. Gel lanes were further cut into smaller pieces, crushed, and soaked overnight at 4 °C in LoTET buffer. Libraries were filtered from gel fragments through 0.22-or 0.45-µm cellulose acetate centrifugal filters (Thermo Scientific) and concentrated to ∼500 μl using n-butanol. Finally, they were cleaned up using AMPure XP beads that had been diluted 8× in the bead binding buffer; diluted beads were added to the sample at a 2:1 ratio, and the final product was resuspended in 20 μl LoTET buffer.

### Sequencing

Libraries were sequenced using XLEAP-SBS chemistry. RPE1 hTERT FUCCI cell cycle stage and translation inhibition datasets were sequenced on a NextSeq2000 (Illumina) with 65 cycles for read 1, 55 cycles for read 2, 6 cycles for the i7 index read (plate index), and 10 cycles for the i5 index read (cell index). Mouse intestinal dataset was sequenced on a NovaSeq X (Illumina) at 150 cycles for reads 1 and 2, 6 cycles for the i7 index read (plate index), and 10 cycles for the i5 index read (cell index).

### Reference genomes and annotations

Reference genomes were processed as previously described¹. For RPF alignment and gene quantification, reads were aligned to the human (GENCODE Release 49, GRCh38) and mouse (GENCODE Release M38, GRCm39) reference genomes and annotations. For rRNA-cut analyses, species-specific rRNA references were used. Human rRNA references comprised RNA45SN1, RNA5S1-17, MT-RNR1, and MT-RNR2, whereas mouse rRNA references comprised Rn45s, Rn18s, Rn28s, Rn5.8s, Rn5s, mt-Rnr1, and mt-Rnr2. All rRNA reference sequences were obtained from NCBI RefSeq.

### Read processing, parsing, and quantification

Reads were processed as previously described^1^ with modifications. The data was processed in paired-end mode. Hence, adapter sequences were trimmed from both reads using cutadapt to trim R1 using-a TGGAATTCTCGGGT min_overlap=3 and a custom flag added to trim R2 using the UMI sequence for each read. Trimmed reads were aligned to the reference genome using STAR (v. 2.7.10a).

RPF read parsing and quantification were performed as previously described^1^. For the translation inhibitors dataset, six genes with read pile-ups at only a few positions were excluded from the count tables (ENSG00000160789, ENSG00000133703, ENSG00000183873, ENSG00000214756, ENSG00000133316, and ENST00000437139).

rRNA read parsing and quantification. rRNA reads were aligned to the species-specific rRNA references described above. For each rRNA position in each cell, 5′ cuts were defined as the first nucleotide of an aligned read, whereas 3′ cuts were defined as the nucleotide immediately downstream of the final aligned nucleotide.

### Single-cell RPF and rRNA count tables processing

For each dataset, four count matrices were generated: a matrix of unique counts per protein-coding gene sequence (CDS), matrices of 5′ and 3′ cuts at each rRNA position, and a matrix of *P_cut_* values for each rRNA position. *P_cut_* was calculated as the ratio of 5′ cuts to total coverage at each position. Data modalities sequentially went through quality control. For translation inhibitors dataset, cells were filtered out if the fraction of reads aligning to the CDS was lower than 0.6 and total number of reads per cell, including 5’UTR, CDS, and 3’UTR was lower than 1250 or higher than 50,000. For the cell-cycle and mouse intestinal epithelial datasets, cells were filtered to remove outlier cells that did not have a minimum number of reads aligning to protein-coding genes, or a sufficient fraction of reads aligning to CDS regions. For the cell-cycle data, these empirical thresholds were determined in an MNase-concentration-dependent manner, and for the intestinal epithelial data, these were determined for each plate to account for minor differences in sequencing depth. Cells were further filtered based on their rRNA measurements. rRNA quality control was performed separately for each MNase concentration. We first plotted the ratio of 28S to 18S 5′ cuts against the total number of rRNA 5′ cuts per cell. Cells outside empirically defined bounds for either metric were excluded. Thresholds were selected based on the main cell population. We next filtered cells with atypical *P_cut_* profiles. For each cell, we calculated the Euclidean distance between its *P_cut_* profile and the mean profile of retained cells at the same MNase concentration. These distances were Z-score normalized, and cells with Z-scores greater than 2 were excluded.

Analysis of single-cell RPF counts were performed using Seurat (version 5.5.1) following standard workflows for normalization, dimensionality reduction, and clustering. All clustering, differential expression, and pseudotime analyses were performed using the unique protein-coding counts. For analysis of rRNA in translation inhibitors dataset, only positions with mean 5’ cut number > 10 in CTRL were used. The resulting cell-by-rRNA position *P_cut_*table was log-odds normalized, scaled with regressing out plate effect, followed by dimensionality reduction.

### Differential digestion testing

The significance of differential digestion at each base was assessed using beta-binomial GLM regression, using the number of 5’ cuts (i.e., “successes”), number of observations (i.e., “trials”), and a parameter to account for overdispersion. Specifically, at each rRNA position with mean 5′ cut count > 10 in the CTRL condition, we modeled the number of 5′ cuts out of total coverage as a beta-binomial response and fit two models with vglm (VGAM package, R): a full model with experimental condition as a covariate, and a reduced intercept-only model. Significance was assessed by a likelihood ratio test comparing the two models, and per-condition contrasts (PURO vs CTRL, HARR vs CTRL) were performed independently. P-values were adjusted across positions using the Benjamini–Hochberg procedure (FDR). Effect size was reported as the logit difference between conditions, which corresponds to the natural log-odds-ratio scale of the beta-binomial model and, unlike log2 fold change, is symmetric and well-defined across the full [0,1] range of proportions.

### Mathematical modeling of rRNA digestion and sequencing noise

A detailed description of the analytical framework underlying identification and deconvolution of ribosomal states, including model assumptions, treatment of technical noise, and derivation of the *η_xz_ = η_yz_* invariant, is provided in the Supplementary Note.

### Clustering of co-varying rRNA positions

For each MNase digestion level, single-cell 5′ and 3′ cut counts were concatenated into a unified feature matrix. Human cell-cycle datasets were analyzed independently at each MNase digestion level, whereas mouse intestinal datasets were analyzed independently for each experimental batch. The normalized variance statistic *η_xx_*and its relative estimation error were estimated by bootstrap resampling. Positions were initially retained if they exhibited sufficient signal, *η_xx_* values within dataset-specific ranges to remove positions with either negligible or atypically large variability, and relative estimation errors below dataset-specific thresholds. Pairwise normalized covariance statistics *η_xy_* were then computed between all retained positions across cells. To remove positions with weak covariance structure, the 15% of positions with the lowest row-wise sums of squared covariance values, Σᵧηₓᵧ², were discarded. The robustness of this statistic was assessed by bootstrap resampling of cells. For each bootstrap replicate, the full (*η_xy_*) matrix was recomputed, the row-wise sums (Σᵧηₓᵧ²) recalculated, and a signal-to-noise ratio (SNR) was defined as the full-data row-wise sum divided by its bootstrap standard deviation. Only positions with SNR greater than 3 were retained.

Retained positions were represented as a graph in which edges connected pairs of positions exhibiting highly correlated covariance profiles and similar normalized variances. Communities of co-varying positions were identified using the Leiden algorithm as implemented in python-igraph and leidenalg, with the resolution parameter selected separately for each dataset. Cluster robustness was assessed by recomputing η after removal of the 1% of cells with the highest total cut signal across the positions in each cluster. thereby testing the sensitivity of clusters to outlier cells. Clusters whose mean η value changed by more than a dataset-specific threshold following trimming, or that contained 20 or fewer positions, were excluded from further analysis. Dataset-specific parameter values and additional details of graph construction, (*η_xy_*) estimation, bootstrap procedures, and cluster filtering are provided in the Supplementary Note.

### Inference of ribosomal states

For each cell, cuts were summed across positions within each retained cluster to generate a cell-by-cluster count matrix. Prior to non-negative matrix factorization (NMF), cluster counts were normalized by the total abundance of each cluster across all cells and subsequently normalized within each cell. NMF was applied using the Nimfa implementation of the Brunet algorithm. A total of 201 random initializations were performed, and the solution with the lowest residual sum of squares was retained. The resulting factors were rescaled to remove the inherent scale non-uniqueness of NMF, and state fractions were computed from the rescaled state-abundance estimates. Additional details of matrix normalization and scaling are provided in the Supplementary Note.

### Ribosome structure analysis and visualization

The mouse and human ribosome structures were accessed from Protein Data Bank (PDB) 7CPU^32^ and 6QZP^33^, respectively. Contact number for a target nucleotide was quantified with minor modifications^15,16^ as total number nucleotides or amino acids having at least one heavy atom within a given radius from the target nucleotide P-atom (see figure cations for radii). Signature cuts and DDPs were visualized on ribosome structures using PyMOL. Positions within the 45S rRNA reference were converted to mature 18S, 5.8S and 28S rRNA coordinates using annotated rRNA boundaries. Since the 28S rRNA sequence used for read alignment differs slightly from that present in the cryo-EM structures, 28S coordinates were translated by pairwise sequence alignment before structural mapping. Positions corresponding to pre-rRNA regions outside mature rRNAs were excluded from structural visualization. Figure generation was performed using PyMOL (https://www.pymol.org/).

### Secondary-structure visualization of rRNA positions

Signature cuts were visualized on the mouse 28S rRNA secondary-structure model obtained from RNAcentral (URS00026FBA82). Positions within the 45S rRNA reference were converted to mature 28S rRNA coordinates using annotated rRNA boundaries. Because the 28S rRNA sequence used for read alignment differs slightly from that represented in the secondary-structure model, 28S coordinates were subsequently mapped onto the secondary-structure model by local sequence matching. Mapped positions were displayed as colored circles overlaid on the secondary structure.

### Cell cycle ordering by ergodic rate analysis

Cell cycle dynamics were extracted from static population snapshots using Ergodic Rate Analysis (ERA), following the previously published method^30^. First, interphase cells were filtered based on both their sort population and cluster annotations. Mitotic cells were filtered by cluster annotations. Only cells within the defined range of MNase concentrations ([0.39 to 25] U MNase) were used. The cell cycle trajectory was identified by following the ridge of maximum density through 2D FUCCI space using an automated path-finding algorithm with a search radius of 20 pixels and a step size of 5 pixels. The resulting path was smoothed with a window size of 5, and all cells were projected onto this 1D trajectory using nearest-segment projection, creating a 1D density profile with 200 bins. The ERA transformation converts this static density distribution into dynamic cell cycle progression rates using the relationship *w*(*l*) = (2 − *F*(*l*))/*f*(*l*), where *w*(*l*) is the progression rate at position *l*, *F*(*l*) is the cumulative distribution function, and *f*(*l*) is the probability density function. Integration of the inverse rate profile yielded scaled pseudotime values for interphase and mitotic cells separately.

### Mega-SEC

Separation of ribosomal states was performed by Mega-SEC as described before^28,29^ using an Agilent Bio SEC-5 2,000 Å column and ӒKTA pure™ 25 (Cytiva) fast protein liquid chromatography setup equipped with 1/16” outer diameter (OD) PEEK tubing (0.25 mm inner diameter (ID), Cytiva) and with a 0.36 mm OD silica tubing (0.15 mm ID, Upchurch Scientific) contained in a 1/16’’ OD tubing sleeve (NanoTight™, Upchurch Scientific) for the last 6.2 cm before fractionation to enable small volume droplets (8-9 μL). The SEC column was equilibrated with five column volumes (CV) of MQ followed by three CV of filtered SEC buffer (20 mM HEPES-NaOH pH7.5, 60 mM NaCl, 10 mM MgCl₂, 0.3% CHAPS, 2.5 mM DTT) at 5°C. After each Mega-SEC experiment, the equipment was washed as follows: system wash was performed on the ӒKTA pure™ and the injection port was flushed with MQ and SEC buffer. Loop, syringe and injection adapter was rinsed in 70% EtOH, MQ followed by SEC buffer, and the column was re-equilibrated with three CV SEC buffer. 200 μL lysate containing 12.6-12.9 μg RNA was injected into the column. Absorbance profiles were recorded using a U9-M with 1 cm pathlength, the multi-wavelength UV/Vis monitor, at 230, 260, and 280 nm. 250 μL fractions were collected at a 0.2 mL min^-1^ flow rate. Data was collected for 4 independent experiments, however, one experiment was excluded from further analysis because of a pipetting mistake.

Raw absorbance spectra were analyzed by normalizing peak intensity to the area under the curve. Gaussian curve (40S peak) or exponentially-modified Gaussian curves (other states) were fitted to the raw or area under the curve normalized peaks in a sequential manner (polysomes, 60S, monosomes, and 40S), with every new curve being fitted to the residuals of the previous fitting.

### Protein mass-spectrometry

For protein mass spectrometry, individual fractions were either pooled (polysomes and monosomes) or processed separately (60S, 40S, and subribosomal fractions) to correct for differences in protein concentration between fractions and aiming at 20-25 μg protein per sample based on absorbance spectra at 260 and 280 nm. The volume of the fractions was reduced to 50 µl in the SpeedVac after which 5 µl 5% SDS, 100 mM Tris(2-carboxyethyl)phosphine hydrochloride (TCEP), 400 mM 2-chloro-acetamide (CAA) was added to denature the proteins and alkylate the cysteines. Proteins were cleaned up with the SP3 protocol^34^ and further digested overnight at 37 °C on heater-shaker with 0.5 µg Trypsin (Whortingthon) and 0.2 µg LysC (Santa Cruz) in 100 mM HEPES pH 8.0, 10 mM CaCl_2_. After acidification with 2% formic acid (FA), peptides were separated from the beads and loaded on C-18 stage tips (Affinisep). Following elution from the stage tips, acetonitrile was removed using a SpeedVac, and the remaining peptide solution was diluted with buffer A (0.1% FA in water) before loading. Peptides were separated after trapping on a pre-column on a 25 cm in-house-made pulled emitter fused silica column (75 µm ID, Polymicro) packed with 1.9 µm aquapur gold C-18 material (dr. Maisch) using a 1 h gradient (5% to 80% acetonitrile/0.1% FA), delivered by an Vanquish Neo HPLC (Thermo), and electro-sprayed directly into a Orbitrap Astral Mass Spectrometer (Thermo Scientific). The latter was set in data independent mode with a cycle time of 0.6 s for both Faims CV settings (-45V and -65V), in which the full scan was performed at a resolution of 240,000. The precursor mass range for both CVs was 380-980 Th in which the peptides, from an isolation window of 2 Th, were fragmented with a normalized collision energy of 25% and an AGC of 500%.

### Spectronaut Data Search

The raw mass spectrometry data was searched using Spectronaut Biognosys (version 20.1.250624.92449) with the directDIA workflow. The reference proteome of Homo sapiens, downloaded in September 2024 from UniProt, was used for the protein identification, a set of common contaminant sequences was also included. Enzymatic digestion was specified as trypsin/P, allowing up to two missed cleavages. Carbamidomethylation was set as a fixed modification, while N-terminal protein acetylation and methionine oxidation were treated as variable modifications. Default (BGS Factory) parameters were applied for the search. Protein quantification was based on MS1 signal intensities, using the MaxLFQ algorithm for label-free quantification, cross-run normalization was enabled and performed automatically, and a false discovery rate threshold of 1% (q-value ≤ 0.01) was applied for both peptide and protein levels. Protein group iBAQ normalized intensities were extracted from the report and calculated as the sum of all identified quantities of the elution groups divided by the number of theoretical specific peptides without any missed cleavages. Data was collected for four independent experiments, however, one experiment was omitted from further analysis due to a pipetting mistake during Mega-SEC. Proteins identified by only one peptide were excluded from the analysis. Proteins identified with posterior error probability ≤ 0.01 in 3 replicates in at least one condition-fraction group were retained for downstream analysis. Principal component analysis (PCA) was performed on proteins identified across all samples.

## Acknowledgements

We thank members of the van Oudenaarden laboratory for helpful discussions; A. Didriksen for his assistance with size-exclusion chromatography experiments; Hubrecht Sorting Facility and the Utrecht Sequencing Facility, subsidized by the University Medical Center Utrecht, the Hubrecht Institute, Utrecht University and The Netherlands X-omics Initiative (NWO project 184.034.019). This work was supported by a European Research Council Advanced grant (ERC-AdG grant no. 101053581-scTranslatomics). This work is part of the Oncode Institute, which is partly financed by the Dutch Cancer Society. H.R.V., P.S.A. and R.M.v.E. are supported by the Oncode Accelerator Project, the Dutch National Growth Fund (NGFOP2201).

## Author information

These authors contributed equally: Euan Joly-Smith, Michael VanInsberghe, Kseniia Sarieva, Eugenio Marinelli.

## Contributions

E.J., M.V., K.S., E.M., and A.v.O. conceived and designed the project. M.V. developed the experimental protocols for the RPE1-FUCCI cell MNase titration and mouse experiments and performed the experiments. K.S. developed the translation inhibition and Mega-SEC experimental protocol and performed the experiments. E.J and E.M. conceived the state deconvolution strategy. E.J. developed the state deconvolution computational pipeline. E.J., M.V., K.S., E.M., and A.v.O. analysed and interpreted the data. E.J., K.S., E.M. and A.v.O. wrote the manuscript with feedback from M.V. A.A.-R. and H.C. provided research materials. R.M.v.E. and H.R.V. performed protein mass spectrometry. P.S.A. and K.S. analyzed protein mass spectrometry data. E.J., M.V., K.S., E.M., and A.v.O. discussed the results and contributed to the interpretation of the findings. All authors approved the final manuscript.

## Corresponding authors

Correspondence to Alexander van Oudenaarden.

## Ethics declarations

All mouse experiments were conducted under the project license AVD8010020151 granted by the Dier Experiment Commissie/Animal Experimentation Committee (DEC) or Central Committee Animal Experimentation (CCD) of the Dutch government and approved by the Hubrecht Institute Animal Welfare Body (IvD).

