## Extended Figures for "Single-Cell Inference of Structural States Of Ribosomes"

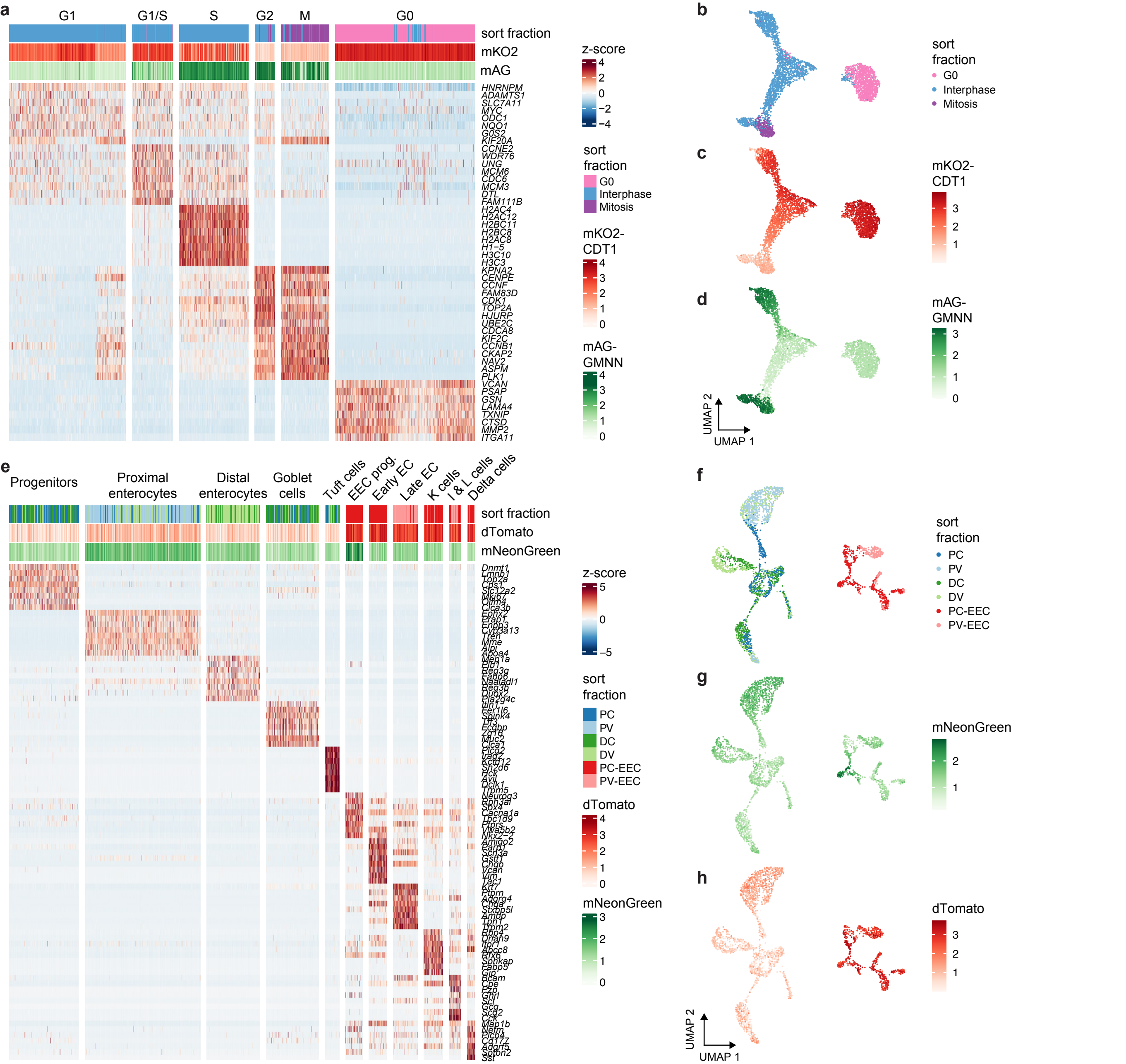
a
G1
G1/S
S
G2
M
G0
sort fraction
mKO2
mAG
HNRNPM
ADAMTS1
SLC7A11
MYC
ODC1
NQO1
G0S2
KIF20A
CCNE2
WDR76
UNG
MCM6
CDC6
MCM3
DTL
FAM111B
H2AC4
H2AC12
H2BC11
H2BC8
H2AC8
H1-5
H3C10
H3C3
KPNA2
CENPF
CCNF
FAM83D
CDK1
TOP2A
HJURP
UBE2C
CDCA8
KIF2C
CCNB1
CKAP2
NAV2
ASPM
PLK1
VCAN
PSAP
GSN
LAMA4
TXNIP
CTSD
MMP2
ITGA11
z-score
4
2
0
−2
−4
sort fraction
G0
Interphase
Mitosis
mKO2-CDT1
4
3
2
1
0
mAG-GMNN
4
3
2
1
0
b
sort fraction
G0
Interphase
Mitosis
c
mKO2-CDT1
3
2
1
d
mAG-GMNN
3
2
1
0
UMAP 2
UMAP 1
e
Progenitors
Proximal enterocytes
Distal enterocytes
Goblet cells
Tuft cells
EEC prog.
Early EC
Late EC
K cells
I & L cells
Delta cells
sort fraction
dTomato
mNeonGreen
z-score
5
0
−5
sort fraction
PC
PV
DC
DV
PC-EEC
PV-EEC
dTomato
4
3
2
1
0
mNeonGreen
3
2
1
0
f
sort fraction
PC
PV
DC
DV
PC-EEC
PV-EEC
g
mNeonGreen
2
1
h
dTomato
3
2
1
UMAP 2
UMAP 1

**Extended Data Figure 1. Marker genes used to identify cell types. a**, Heatmap showing example marker genes differentially expressed between cell-cycle stages. Cells (columns) are annotated with the cell fraction from which they were sorted (sort fraction) and the fluorescence of the monomeric Kusabira-Orange 2 (mKO2)–CDT1 and monomeric Azami-Green (mAG)–GMNN FUCCI markers collected during index sorting. **b-d**, UMAPs (n = 3678 cells) illustrating **b**, the sort fraction, **c**, fluorescence of the mKO2-CDT1 and **d**, mAG-GMMN FUCCI markers. **e**, Heatmap showing example marker genes differentially expressed between mouse intestinal epithelial cell types. Cells (columns) are annotated with the cell fraction from which they were sorted and the fluorescence of the dTomato and mNeonGreen reporters from the Neurog3Chrono mouse[31] collected during index sorting. PC: proximal crypt; PV: proximal villi; DC: distal crypt; DV: distal villi; EEC: enteroendocrine. **f-h**, UMAPs (n = 2659 cells) illustrating **f**, the cell fraction, **g,** fluorescence of the destabilized mNeonGreen and **h**, and dTomato reporters.

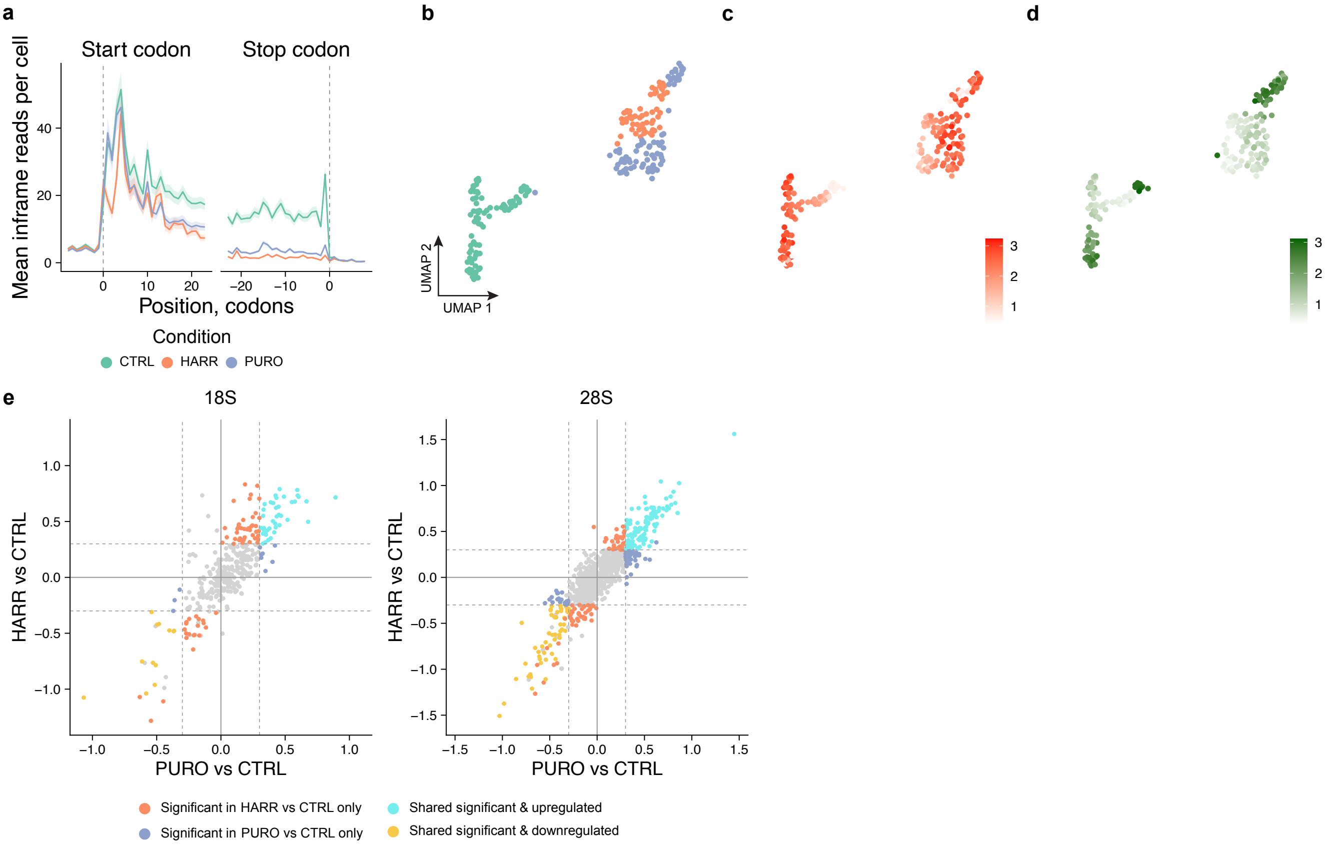
a
Start codon
Stop codon
Mean inframe reads per cell
Position, codons
Condition
CTRL
HARR
PURO
b
UMAP 1
UMAP 2
c
d
e
18S
28S
HARR vs CTRL
PURO vs CTRL
Significant in HARR vs CTRL only
Significant in PURO vs CTRL only
Shared significant & upregulated
Shared significant & downregulated

**Extended Data Figure 2. Translation inhibitors alter ribosomal state distribution and cellular translatome. a,** Metagene plot illustrating average predicted P-site count per cell along regions aligned around the start codon (left) and the stop codon (right). **b-d,** UMAPs (n = 209 cells) of RPE1-FUCCI cells based on RPF counts per CDS illustrating treatment conditions (**b**) monomeric Kusabira-Orange 2 (mKO2)–CDT1 marker fluorescence (**c**), and monomeric Azami-Green (mAG)–GMNN FUCCI marker fluorescence (**d**). **e,** Differential rRNA digestion analysis between two comparisons (PURO vs CTRL, HARR vs CTRL) for 18S and 28S separately. Differentially digested positions are highlighted (abs(logit difference) > 0.3, adjusted p-value < 0.05, beta-binomial test, Benjamini-Hocheberg correction).

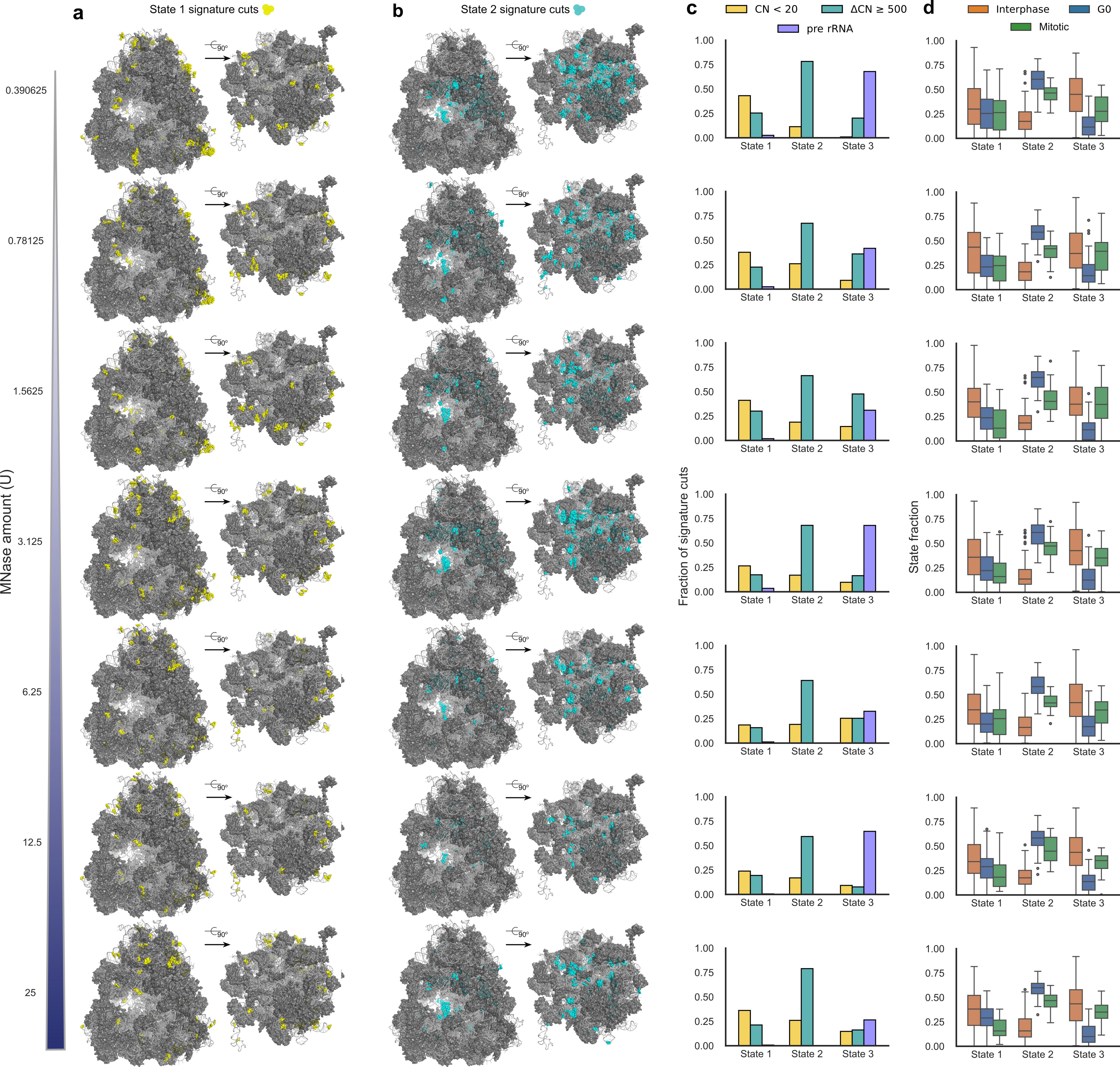
a
State 1 signature cuts
b
State 2 signature cuts
c
CN < 20
ΔCN ≥ 500
pre rRNA
d
Interphase
G0
Mitotic
MNase amount (U)
0.390625
0.78125
1.5625
3.125
6.25
12.5
25
90°
Fraction of signature cuts
State fraction
State 1
State 2
State 3

**Extended Data Figure 3. Ribosomal states are reproduced across multiple MNase digestion levels. a,** State 1 signature cuts highlighted on cryo-EM models of the human ribosome (PDB: 6QZP) across increasing MNase digestion levels (0.195–25 U), showing enrichment at outer surface regions. **b,** State 2 signature cuts highlighted on cryo-EM models of the human ribosome, showing enrichment at the intersubunit interface across multiple MNase concentrations. **c,** Fraction of signature cuts for each state with CN < 20 (with 12 Å radius), ΔCN ≥ 500 (with 100 Å radius), or mapping to pre-rRNA across multiple MNase concentrations. State 1 consistently exhibits strong enrichment for cuts with CN < 20, corresponding to regions near the ribosomal outer surface. State 2 consistently exhibits enrichment for cuts with ΔCN ≥ 500, corresponding to regions near the intersubunit interface. State 3 consistently exhibits enrichment in pre-rRNA. **d,** Distribution of NMF state fractions across cell-cycle populations at increasing MNase digestion levels. Box plots show the fraction of reads assigned to States 1–3 in Interphase, G0, and Mitotic cells for each MNase amount. Boxes indicate the interquartile range (IQR), center lines denote medians, and whiskers extend to 1.5 × IQR.

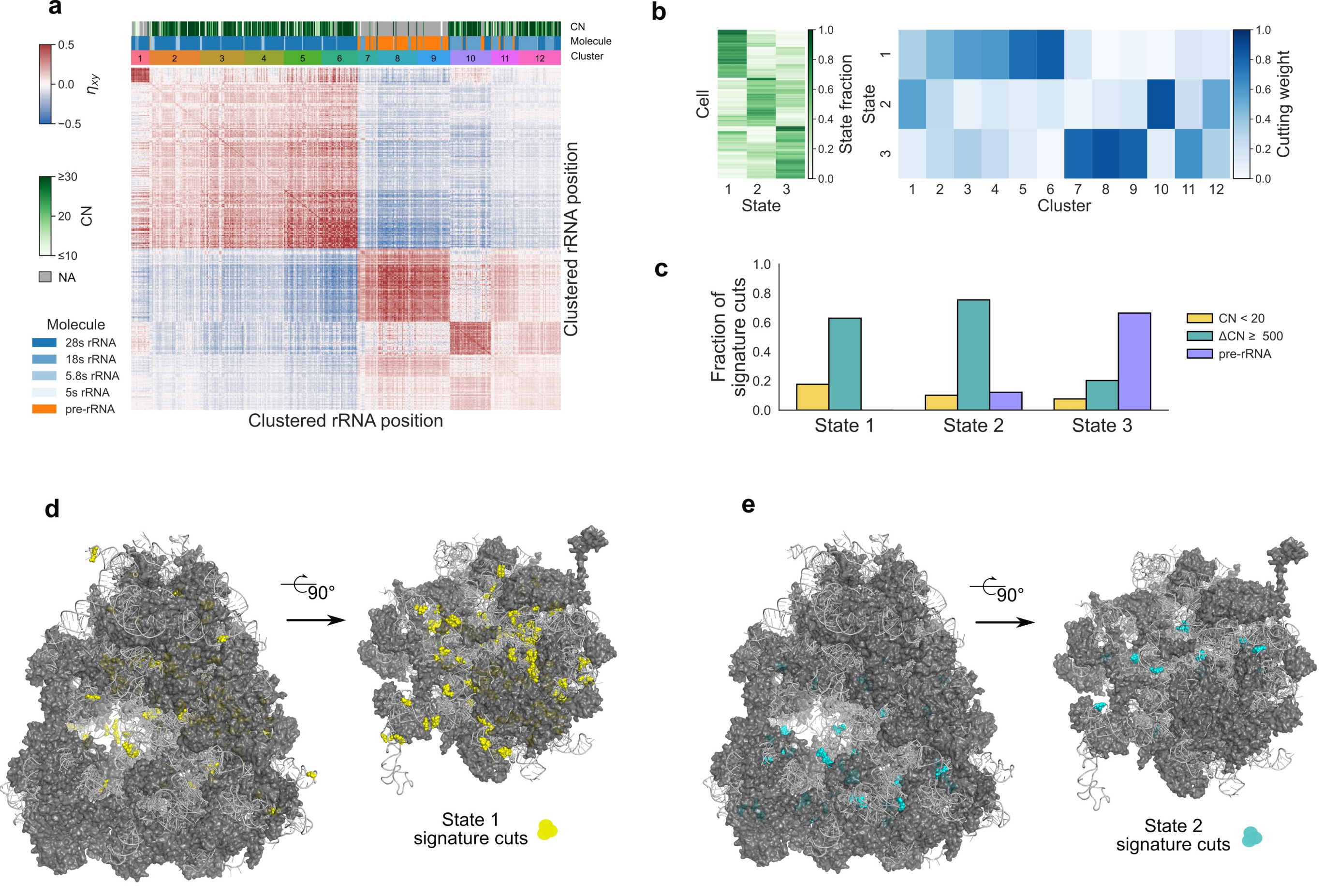
a
CN
Molecule
Cluster
1 2 3 4 5 6 7 8 9 10 11 12
0.5
0.0
−0.5
$\eta_{xy}$
≥30
20
≤10
CN
NA
Molecule
28s rRNA
18s rRNA
5.8s rRNA
5s rRNA
pre-rRNA
Clustered rRNA position
Clustered rRNA position
b
Cell
State
State fraction
State
Cluster
Cutting weight
c
Fraction of signature cuts
State 1
State 2
State 3
CN < 20
ΔCN ≥ 500
pre-rRNA
d
90°
State 1 signature cuts
e
90°
State 2 signature cuts

**Extended Data Figure 4. NMF states are not readily interpretable at low MNase digestion levels. a,** Heatmap of measured $\eta_{xy}$ values at MNase amount 0.024 U, illustrating clustered rRNA positions at low MNase digestion. Three color bars indicate the CN (with 12 Å radius) for each position, molecule identity (pre-rRNA, 18S, 28S, or 5.8S rRNA), and cluster assignment. Unlike Fig. 3b, clustering is not associated with low-CN regions. **b,** NMF output matrices showing per-cell state fractions (left) and cutting weights (right). **c,** Fraction of signature cuts for each state with CN < 20 (with 12 Å radius), ΔCN ≥ 500 (with 100 Å radius), or mapping to pre-rRNA. In contrast to the states identified at higher MNase digestion levels, none of the three states exhibit strong enrichment for signature cuts with CN < 20. **d,** Cryo-EM model of the human ribosome with State 1 signature cuts highlighted. Unlike the surface-enriched pattern observed at higher MNase digestion levels, signature cuts are enriched at the interface at the intersubunit interface. **e,** State 2 signature cuts are enriched at the interface.

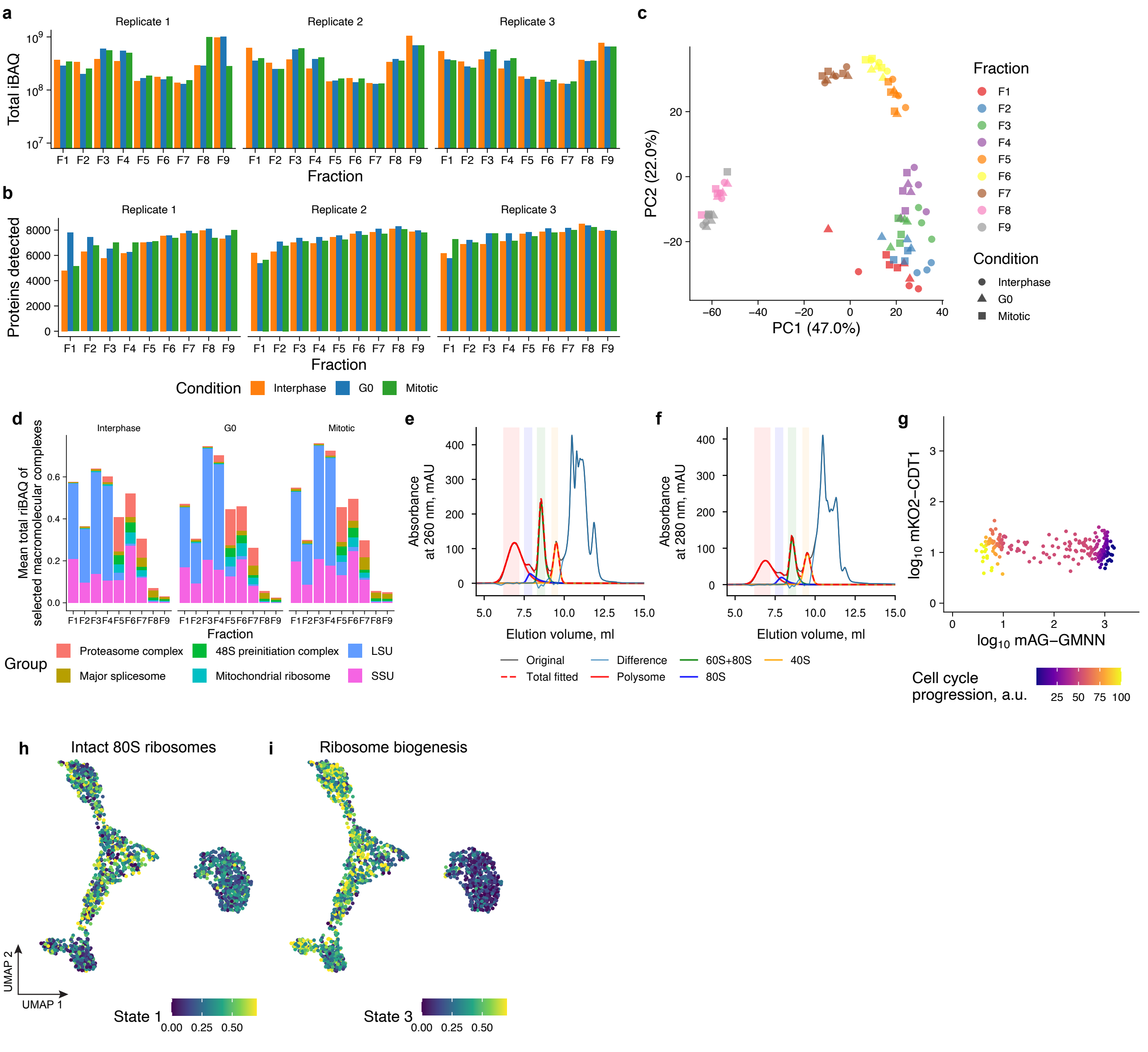
a
Replicate 1
Replicate 2
Replicate 3
Total iBAQ
Fraction
b
Proteins detected
Fraction
Condition Interphase G0 Mitotic
c
PC1 (47.0%)
PC2 (22.0%)
Fraction
F1
F2
F3
F4
F5
F6
F7
F8
F9
Condition
Interphase
G0
Mitotic
d
Interphase
G0
Mitotic
Mean total riBAQ of selected macromolecular complexes
Fraction
Group
Proteasome complex
48S preinitiation complex
LSU
Major splicesome
Mitochondrial ribosome
SSU
e
Absorbance at 260 nm, mAU
Elution volume, ml
f
Absorbance at 280 nm, mAU
Elution volume, ml
Original
Difference
60S+80S
40S
Total fitted
Polysome
80S
g
log10 mKO2−CDT1
log10 mAG−GMNN
Cell cycle progression, a.u.
h
Intact 80S ribosomes
i
Ribosome biogenesis
UMAP 1
UMAP 2
State 1
State 3

**Extended Data Figure 5. Ribosomal state distributions and associated proteins vary between cell cycle stages. a,b,** total iBAQ (**a**) and number of proteins (**b**) per sample (experiment x cell cycle stage x condition x fraction) after quality control. **c**, PCA using the iBAQ of overlapping proteins (2,397). Different colors denote fractions (consistent with Fig. 4c); shapes denote cell cycle stages. **d**, Summarised riBAQ of proteins belonging to different macromolecular complexes across fractions. Mean values across three experiments are shown. **e,f,** Representative examples of exponentially modified Gaussian (polysome, 60S+80S, and 80S peaks) and Gaussian (40S) sequential curve fitting to absorbance profiles at 260 (**e**) and 280 (**f**) nm. Colors of lines represent raw absorbance measurement (Original), individual curves, total fitted, and total difference between measurement and fitting. Colored stripes show the ranges for maximum estimation for individual curves. **g,** Scatter plots of the FUCCI markers with colors denoting time along cell cycle progression as calculated by ERA for mitotic cells. **h,i,** UMAP of RPE1-FUCCI cells based on RPFs. Cells treated with multiple MNase concentrations are shown. Color represents the fraction of state 1 (**h**) and state 3 (**i**) as assessed by NMF.

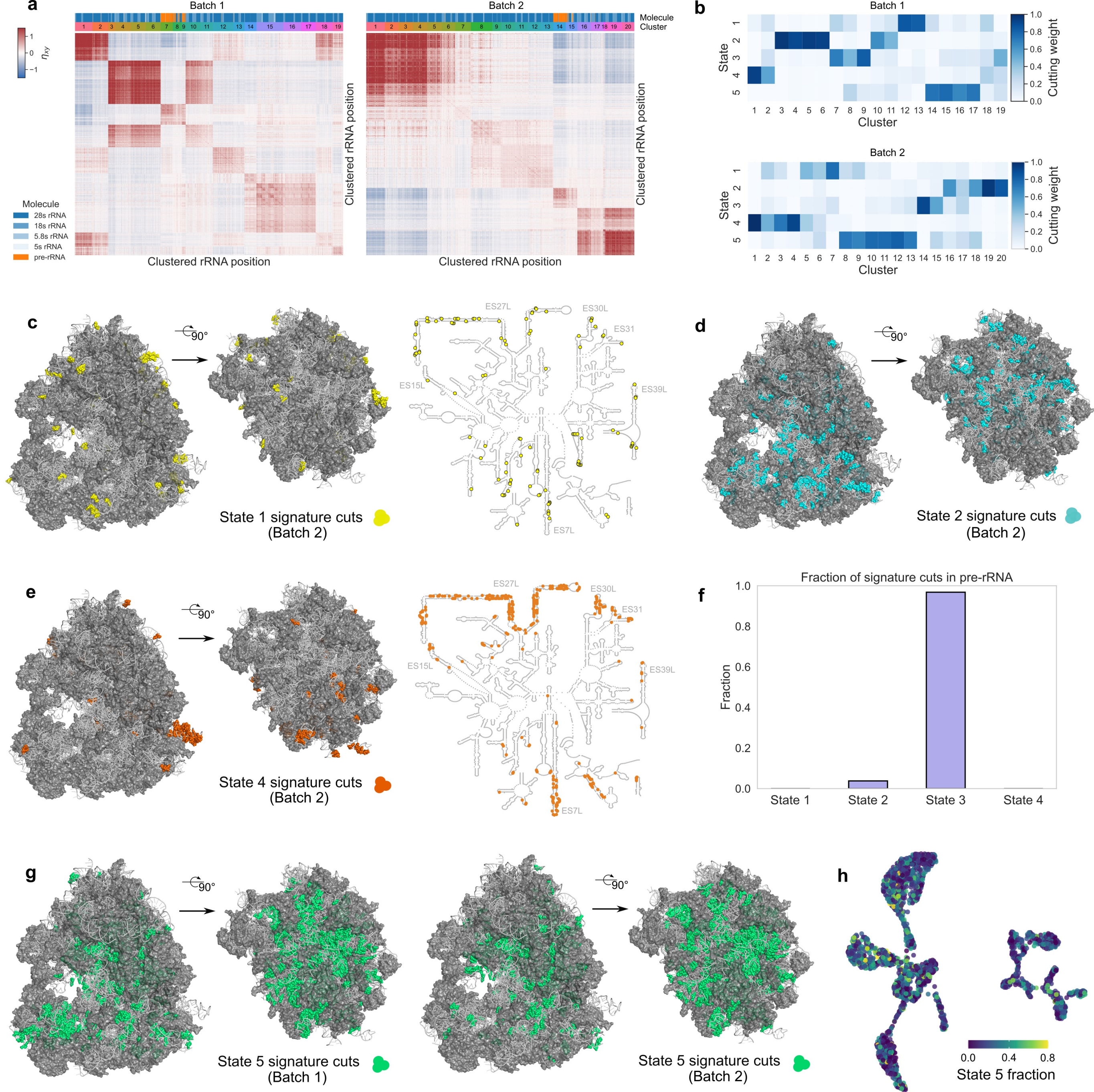
a
Batch 1
Batch 2
Molecule
Cluster
$\eta_{xy}$
Clustered rRNA position
Clustered rRNA position
Clustered rRNA position
Clustered rRNA position
Molecule
28s rRNA
18s rRNA
5.8s rRNA
5s rRNA
pre-rRNA
b
Batch 1
Batch 2
State
Cluster
Cutting weight
c
90°
ES27L
ES30L
ES31
ES15L
ES39L
ES7L
State 1 signature cuts
(Batch 2)
d
90°
State 2 signature cuts
(Batch 2)
e
90°
ES27L
ES30L
ES31
ES15L
ES39L
ES7L
State 4 signature cuts
(Batch 2)
f
Fraction of signature cuts in pre-rRNA
Fraction
State 1
State 2
State 3
State 4
g
90°
State 5 signature cuts
(Batch 1)
90°
State 5 signature cuts
(Batch 2)
h
State 5 fraction

**Extended Data Figure 6. NMF states from murine intestine are reproducible across two independent batches. a,** Heatmap of measured $\eta_{xy}$ values, illustrating clustered rRNA positions for two batches with slightly different cell-type abundances. Two color bars indicate molecule identity (pre-rRNA, 18S, 28S, or 5.8S rRNA) and cluster assignment. **b,** NMF output matrices showing cutting weights for each batch. Cutting weights and signature cuts vary between batches, as MNase cutting probabilities may depend on experimental conditions following cell lysis. **c,** Cryo-EM model of the mouse ribosome with State 1 signature cuts from batch 2 highlighted. Signature cuts are enriched at the ribosomal outer surface. **d,** State 2 signature cuts from batch 2 are enriched near the intersubunit interface. **e,** State 4 signature cuts from batch 2. Cuts are enriched within ribosomal expansion segments. **f,** Fraction of signature cuts from each state localized to pre-rRNA in batch 2. The majority of State 3 signature cuts map to pre-rRNA regions. Signature cuts from batch 1 are shown in Fig. 1a–d. **g,** State 5 signature cuts from batch 1 (left) and batch 2 (right). These positions are distributed throughout the ribosome, including regions on the outer surface. Based on this pattern, no biological interpretation was assigned to this state. Running NMF with fewer than five states results in the loss of State 1, which is characterized by outer-surface-enriched signature cuts. **h,** UMAP embedding of intestinal cells derived from ribosome-protected fragments (RPFs) (Fig. 1d), overlaid with the inferred fraction of ribosomal NMF State 5 from two independent batches (see Methods for the batch integration procedure). No strong cell-type-specific enrichment pattern is observed.
