## Supplementary Note for "Single-Cell Inference of Structural States Of Ribosomes"

### Supplementary Information

Euan Joly-Smith, Michael VanInsberghe, Kseniia Sarieva, Eugenio Marinelli,
Robert M. van Es, Paula Sobrevals Alcaraz, Harmjan R. Vos, Amanda Andersson-Rolf, Hans Clevers,
Alexander van Oudenaarden

**Contents**

I. Theoretical framework 1
A. MNase digestion and fragment detection 1
B. 5p-start counts from a single ribosomal state 1
C. Total 5p-start counts across ribosomal states 2
D. Derivation of the co-variability invariant 2
E. Using cut clusters to quantify ribosomal state mixtures 5
1. Clustering positions by shared co-variability structure 5
2. Reduction of sampling noise by summing within clusters 5
3. Summed cluster counts provide linear readouts of state abundances 6
F. Inferring relative ribosomal state abundances via non-negative matrix factorization 6
G. Identifying ribosomal states using signature cuts 8
H. Cross-experiment comparison of ribosomal state composition 9
I. Incorporating 3p-end information 9

II. Computational workflow for state deconvolution 10
A. Computing single-cell cutting vectors 10
B. Filtering positions for robust covariance estimation 11
C. Computing the normalized covariance matrix $\eta$ 12
D. Clustering the $\eta$ matrix 12
E. Non-negative matrix factorization of summed cluster counts 13
1. Scaling input matrix 13
2. Non-negative matrix factorization procedure 13

III. Mega-SEC Analysis 14
A. Quantification of total ribosomes 14
B. De-mixing of the 60S fraction 14
C. Fraction of free 60S subunits 15

References 15

### I. Theoretical framework

#### A. MNase digestion and fragment detection

MNase digestion is an enzymatic process in which micrococcal nuclease (MNase) cleaves rRNA molecules probabilistically at specific positions along their sequence, generating a collection of fragments of varying lengths. Each rRNA molecule can exist in one of several ribosomal states $i = 1, \ldots, N_{\text{states}}$. The structural differences between states lead to different MNase cutting probabilities.

Specifically, for a rRNA molecule in state $i$, let $\pi_i(x, l, m)$ denote the probability that MNase exposure at concentration $m$ produces a fragment whose 5p start is at reference position $x$ and whose length is $l$. For each rRNA molecule, there can be up to one fragment that starts at $x$, thus $\pi_{i,\text{null}}(x, m) + \sum_l \pi_i(x, l, m) = 1$ where $\pi_{i,\text{null}}(m)$ is the probability that there is no cut at position $x$ on this particular molecule following the digestion step.

After digestion, the contents of the cell are sequenced. During this process, each fragment present must successfully be processed through library preparation, amplification, sequencing, and computational mapping. This introduces a probabilistic detection step that adds measurement noise to the observed data. In particular, if a fragment with start at $x$ with length $l$ is produced during digestion, it is then detected with probability $\beta(x, l)$, an unspecified probability which encompasses length-dependent sequencing biases (such as the inability to map very short reads), GC-content effects, and sequencing efficiency.

The probability that a single rRNA molecule in state $i$ yields a *detected* fragment beginning at position $x$ is

$$p_i(x, m) = \sum_l \beta(x, l)\, \pi_i(x, l, m), \tag{1}$$

which we call the *effective start probability.* It summarizes the combined effect of MNase cutting and detection across all possible fragment lengths.

#### B. 5p-start counts from a single ribosomal state

Consider a cell that contains $n_i$ rRNA molecules in state $i$. Assuming that MNase digests each such molecule independently, and each fragment that exists after digestion is detected independently according to $\beta$, it follows that the number of detected fragments starting at $x$ from state $i$ is binomially distributed:

$$n_{5\text{p},i}(x, m) \sim \text{Binomial}(n_i,\ p_i(x, m))\,. \tag{2}$$

In addition, we make two small-probability approximations that will be used later in the covariance derivation. First, we assume that the effective start probabilities $p_i(x, m)$ are small. This assumption is supported by the data. A single mammalian cell contains on the order of $10^6$-$10^7$ ribosomes [1], yet the number of detected 5p starts at any given position $x$ ranges broadly from $\sim 1$ to $\sim 10^3$ per cell. Even at the upper end of this range,

$$p_i(x, m) \sim \frac{10^3}{10^3} \approx 10^{-3},$$

and for most positions $p_i(x, m)$ is in the range $10^{-6}$–$10^{-5}$. Under these conditions, the probability that a single rRNA molecule contributes multiple detected 5p starts is negligible.

Second, we assume that the count at any individual position contributes only weakly to the total number of detected 5p starts,

$$N_{\text{tot}} = \sum_u n_{5\text{p}}(u, m).$$

Empirically, the normalized frequency at any one position is small, typically on the order of

$$f_{5\text{p}}(x) = \frac{n_{5\text{p}}(x, m)}{N_{\text{tot}}} \sim 10^{-3}.$$

We therefore treat $N_{\text{tot}}$ as approximately independent of the count at any individual position when conditioning on the state abundances $\{n_i\}$ and effective start probabilities $\{p_i\}$. This approximation is used in the derivation of the covariance invariant.

##### C. Total 5p-start counts across ribosomal states

Sequencing a fragment cannot distinguish the ribosomal state from which that fragment originated. The observed 5p start count at position $x$ in a single cell is therefore the sum of contributions over all states:

$$n_{5\mathrm{p}}(x,m) = \sum_{i=1}^{N_{\mathrm{states}}} \mathrm{Bin}(n_i,\ p_i(x,m))\,. \tag{3}$$

When analyzing multiple cells, both the state abundances $n_i$ and the detection biases implicit in $p_i(x,m)$ vary from cell to cell. In particular, $n_i$ reflect biological variability in the abundances of ribosomal states, while effective start probabilities $p_i(x,m) = \sum_l \beta(x,l)\,\pi_i(x,l,m)$ vary through technical noise in $\beta(x,l)$, which differs between wells due to sequencing efficiency. Thus, in a dataset of a population of cells, the collection $\{n_{5\mathrm{p}}^{(c)}(x,m)\}_{c=1}^{N_{\mathrm{cells}}}$ is generated by drawing repeatedly these unspecified random variables for each cell (indexed by $c$).

This formulation provides the generative model underlying the inference of the ribosomal states described in Fig. 3 of the main text.

##### D. Derivation of the co-variability invariant

We define the normalized 5p-start frequency at position $x$ as

$$f_{5\mathrm{p}}(x) = \frac{n_{5\mathrm{p}}(x)}{N_{\mathrm{tot}}}, \quad \text{where} \quad N_{\mathrm{tot}} = \sum_y n_{5\mathrm{p}}(y).$$

Cuts at positions on the rRNA that share similar effective cutting profiles across ribosomal states fluctuate together across cells. If two positions $x$ and $y$ have proportional start probabilities in every ribosomal state, then any cell-to-cell change in the underlying state mixture scales their expected 5p-start frequencies in exactly the same way. As a result, $f_{5\mathrm{p}}(x)$ and $f_{5\mathrm{p}}(y)$ exhibit the same pattern of covariance with all other positions.

However, the observed *variance* of each $f_{5\mathrm{p}}(x)$ also contains an additional contribution from sampling noise, arising from a finite number of detected fragments. After removing this sampling noise component, the remaining variability reflects cell-to-cell differences in ribosomal state composition and sequencing efficiency.

The theorem below formalizes these statements: positions with proportional effective start probabilities have identical normalized co-variability profiles, and their cross-covariance equals the diagonal covariance minus the sampling noise term.

**Theorem 1.** *Let $x$ and $y$ be two reference positions on the rRNA. If the effective start probabilities at $x$ and $y$ are relatively equal across all ribosomal states at a given MNase concentration (i.e., $p_i(x,m) \propto p_i(y,m)$ for all $i$ with fixed proportionality factor), then under the approximations introduced in Sec. I B:*

$$\eta_{xz} = \eta_{yz} \text{ for all other reference positions } z,$$

*where $\eta_{xz} = \mathrm{Cov}\left[f_{5\mathrm{p}}(x), f_{5\mathrm{p}}(z)\right]/\langle f_{5\mathrm{p}}(x)\rangle\langle f_{5\mathrm{p}}(z)\rangle$ is the normalized covariance between the frequency of starts taken over the cell population. In addition, under the same conditions, the following holds:*

$$\eta_{xy} = \eta_{xx} - \frac{\langle f_{5\mathrm{p}}(x)/N_{\mathrm{tot}}\rangle}{\langle f_{5\mathrm{p}}(x)\rangle^2} = \eta_{yy} - \frac{\langle f_{5\mathrm{p}}(y)/N_{\mathrm{tot}}\rangle}{\langle f_{5\mathrm{p}}(y)\rangle^2}.$$

*Proof.* We drop the explicit MNase concentration $m$ dependence in the functions below, since all quantities are evaluated at fixed $m$. Let

$$f_x = f_{5\mathrm{p}}(x) = \frac{n_{5\mathrm{p}}(x)}{N_{\mathrm{tot}}}.$$

For a molecule $r$ in ribosomal state $i$, define the indicator variable

$$A_{ir}(x) = \begin{cases} 1, & \text{if molecule } r \text{ in state } i \text{ produces a detected 5p start at } x, \\ 0, & \text{otherwise.} \end{cases}$$

Then

$$n_{5\mathrm{p}}(x) = \sum_i \sum_{r=1}^{n_i} A_{ir}(x), \qquad \mathbb{E}[A_{ir}(x) \mid \{n_i\}, \{p_i\}] = p_i(x).$$

We use the sparsity approximation that any two detected 5p starts arise from distinct rRNA molecules. Equivalently, for distinct positions $x \neq z$,

$$A_{ir}(x)A_{ir}(z) \approx 0.$$

For $x \neq z$, conditioning on $\{n_i\}$ and $\{p_i\}$,

$$\begin{aligned}\mathrm{Cov}[n_{5\mathrm{p}}(x), n_{5\mathrm{p}}(z) \,|\, \{n_i\}, \{p_i\}] &= \sum_i \sum_{r=1}^{n_i} \mathrm{Cov}[A_{ir}(x), A_{ir}(z) \,|\, \{n_i\}, \{p_i\}] \\ &= -\sum_i n_i\, p_i(x)p_i(z),\end{aligned}$$

where cross-molecule covariance terms vanish by independence.

Now assume that positions $x$ and $y$ satisfy

$$p_i(x) = \alpha\, p_i(y), \qquad \forall i,$$

with $\alpha > 0$. Then, for any third position $z \neq x, y$,

$$\mathrm{Cov}[n_{5\mathrm{p}}(x), n_{5\mathrm{p}}(z) \,|\, \{n_i\}, \{p_i\}] = \alpha\ \mathrm{Cov}[n_{5\mathrm{p}}(y), n_{5\mathrm{p}}(z) \,|\, \{n_i\}, \{p_i\}]\,.$$

Since each individual position contributes only a very small fraction of the total detected starts, $N_{\mathrm{tot}}$ is approximately independent of the count at any one position while conditioning on the $n_i$ and $p_i$. Therefore,

$$\mathrm{Cov}(f_x, f_z \mid \{n_i\}, \{p_i\}) = \mathbb{E}\left[\frac{1}{N_{\mathrm{tot}}^2} \,\middle|\, \{n_i\}, \{p_i\}\right] \mathrm{Cov}(n_{5\mathrm{p}}(x), n_{5\mathrm{p}}(z) \mid \{n_i\}, \{p_i\}),$$

and the same prefactor multiplies the corresponding covariance involving $y$. Hence

$$\mathrm{Cov}(f_x, f_z \mid \{n_i\}, \{p_i\}) = \alpha\, \mathrm{Cov}(f_y, f_z \mid \{n_i\}, \{p_i\}). \tag{4}$$

Similarly,

$$\mathbb{E}[n_{5\mathrm{p}}(x) \mid \{n_i\}, \{p_i\}] = \sum_i n_i p_i(x) = \alpha \sum_i n_i p_i(y) = \alpha\, \mathbb{E}[n_{5\mathrm{p}}(y) \mid \{n_i\}, \{p_i\}],$$

and therefore

$$\mathbb{E}[f_x \mid \{n_i\}, \{p_i\}] = \alpha \mathbb{E}[f_y \mid \{n_i\}, \{p_i\}].$$

Applying the law of total covariance together with Eq. (4) gives

$$\mathrm{Cov}(f_x, f_z) = \alpha\ \mathrm{Cov}(f_y, f_z), \qquad z \neq x, y. \tag{5}$$

Similarly, the law of total expectation gives

$$\langle f_x \rangle = \alpha\, \langle f_y \rangle.$$

Substituting into Eq. (5),

$$\eta_{xz} = \frac{\mathrm{Cov}(f_x, f_z)}{\langle f_x \rangle \langle f_z \rangle} = \frac{\alpha\ \mathrm{Cov}(f_y, f_z)}{\alpha\, \langle f_y \rangle \langle f_z \rangle} = \eta_{yz}, \qquad z \neq x, y.$$

We now derive the diagonal relation. By the law of total variance,

$$\mathrm{Var}(f_x) = \mathbb{E}[\mathrm{Var}(f_x \mid \{n_i\}, \{p_i\})] + \mathrm{Var}(\mathbb{E}[f_x \mid \{n_i\}, \{p_i\}])\,. \tag{6}$$

Using the $N_{\text{tot}}$-independence approximation,

$$\mathrm{Var}(f_x \mid \{n_i\},\{p_i\}) = \mathbb{E}\left[\frac{1}{N_{\text{tot}}^2}\,\middle|\,\{n_i\},\{p_i\}\right]\mathrm{Var}(n_{5\mathrm{p}}(x) \mid \{n_i\},\{p_i\}).$$

Since

$$n_{5\mathrm{p}}(x) = \sum_i \mathrm{Binomial}(n_i, p_i(x)),$$

we have

$$\begin{aligned}\mathrm{Var}(n_{5\mathrm{p}}(x) \mid \{n_i\},\{p_i\}) &= \sum_i n_i p_i(x)\big(1 - p_i(x)\big)\\ &\approx \sum_i n_i p_i(x) = \mathbb{E}[n_{5\mathrm{p}}(x) \mid \{n_i\},\{p_i\}],\end{aligned}$$

where the approximation uses $p_i(x) \ll 1$.

Therefore,

$$\mathrm{Var}(f_x \mid \{n_i\},\{p_i\}) \approx \mathbb{E}\left[\frac{f_x}{N_{\text{tot}}}\,\middle|\,\{n_i\},\{p_i\}\right].$$

Substituting into Eq. (6) gives

$$\mathrm{Var}(\mathbb{E}[f_x \mid \{n_i\},\{p_i\}]) \approx \mathrm{Var}(f_x) - \mathbb{E}\left[\frac{f_x}{N_{\text{tot}}}\right]. \tag{7}$$

Since $p_i(x) = \alpha p_i(y)$ for all $i$,

$$\mathbb{E}[f_x \mid \{n_i\},\{p_i\}] = \alpha\,\mathbb{E}[f_y \mid \{n_i\},\{p_i\}],$$

and therefore

$$\mathrm{Var}(\mathbb{E}[f_x \mid \{n_i\},\{p_i\}]) = \alpha\ \mathrm{Cov}(\mathbb{E}[f_x \mid \{n_i\},\{p_i\}], \mathbb{E}[f_y \mid \{n_i\},\{p_i\}]).$$

By the law of total covariance,

$$\mathrm{Cov}(f_x, f_y) = \mathbb{E}[\mathrm{Cov}(f_x, f_y \mid \{n_i\},\{p_i\})] + \mathrm{Cov}(\mathbb{E}[f_x \mid \{n_i\},\{p_i\}], \mathbb{E}[f_y \mid \{n_i\},\{p_i\}]).$$

Now, recall that the conditional sampling covariance of the raw counts satisfies

$$\mathrm{Cov}[n_{5\mathrm{p}}(x), n_{5\mathrm{p}}(y) \,|\, \{n_i\},\{p_i\}] = -\sum_i n_i p_i(x) p_i(y).$$

Although the $n_i$ are large, this term enters the covariance of normalized frequencies after division by $N_{\text{tot}}^2$. Under the small-compositional-effect approximation,

$$\mathrm{Cov}[f_x, f_y \,|\, \{n_i\},\{p_i\}] \approx -\frac{\sum_i n_i p_i(x) p_i(y)}{N_{\text{tot}}^2}.$$

Since $N_{\text{tot}}$ is large ($\sim 10^5$) and $p_i(x), p_i(y) \ll 1$, this conditional sampling covariance is negligible compared with the cell-to-cell covariance term in the law of total covariance relation. Therefore,

$$\mathrm{Cov}(f_x, f_y) \approx \mathrm{Cov}(\mathbb{E}[f_x \mid \{n_i\},\{p_i\}], \mathbb{E}[f_y \mid \{n_i\},\{p_i\}]).$$

Combining with Eq. (7) yields

$$\alpha\ \mathrm{Cov}(f_x, f_y) \approx \mathrm{Var}(f_x) - \mathbb{E}\left[\frac{f_x}{N_{\text{tot}}}\right].$$

Dividing by $\langle f_x \rangle^2$ and using $\langle f_x \rangle = \alpha \langle f_y \rangle$,

$$\eta_{xy} = \eta_{xx} - \frac{\langle f_x / N_{\text{tot}} \rangle}{\langle f_x \rangle^2}.$$

Thus,

$$\eta_{xy} = \eta_{xx} - \frac{\langle f_x / N_{\text{tot}} \rangle}{\langle f_x \rangle^2} = \eta_{yy} - \frac{\langle f_y / N_{\text{tot}} \rangle}{\langle f_y \rangle^2},$$

where the $y$ relation follows by symmetry. □

##### E. Using cut clusters to quantify ribosomal state mixtures

###### 1. *Clustering positions by shared co-variability structure*

As shown in the previous section, any pair of positions $x$ and $y$ that share the same relative cutting profile,

$$p_i(x) \propto p_i(y) \qquad \text{for all ribosomal states } i,$$

exhibit identical co-variability patterns with all other positions. In addition, after subtracting sampling noise, such pairs achieve the largest possible mutual co-variability, consistent with the Cauchy–Schwarz bound $\eta_{xy} \leq \sqrt{\eta_{xx}\,\eta_{yy}}$.

Thus, by clustering the empirically measured $\eta$ matrix into groups of positions that obey the relation established in Theorem 1, we identify sets of positions with matched co-variability signatures. Positions grouped together are then inferred to share a common effective cutting profile across ribosomal states.

However, individual positions can have widely varying 5p-start counts, and many positions exhibit low counts where sampling noise dominates. We therefore sum counts within each cluster to suppress sampling noise and obtain high-SNR measurements of state mixtures.

###### 2. *Reduction of sampling noise by summing within clusters*

Recall the model for measured 5p-start counts:

$$n_{5\mathrm{p}}(x) = \sum_i \mathrm{Bin}(n_i, p_i(x)).$$

Conditioning on $\{n_i\}$ and $\{p_i(x)\}$ the binomial terms are independent and therefore

$$\mathrm{Var}[n_{5\mathrm{p}}(x) \mid \{n_i\}, \{p_i(x)\}] = \sum_i n_i p_i(x)\big(1 - p_i(x)\big) \approx \sum_i n_i p_i(x),$$

where the last step comes from the fact that $p_i(x) \ll 1$. The unconditional variance is obtained by the law of total variance,

$$\mathrm{Var}(n_{5\mathrm{p}}(x)) = \mathbb{E}\left[\sum_i n_i p_i(x)\right] + \mathrm{Var}\left(\sum_i n_i p_i(x)\right),$$

where the first term reflects sampling noise, and the second term cell-to-cell variability in $n_i$ and $p_i(x)$.

The noise-to-signal ratio can be computed using the sampling component of the unconditional variance:

$$\frac{1}{\mathrm{SNR}} = \frac{\sqrt{\mathrm{Var}[n_{5\mathrm{p}}(x)]_{\mathrm{sampling}}}}{\mathbb{E}[n_{5\mathrm{p}}(x)]} = \frac{1}{\sqrt{\mathbb{E}[n_{5\mathrm{p}}(x)]}}$$

which is significant for positions with average count on the order of $\sim 1$ or $\sim 10$.

Let $C$ be a cluster of positions with same relative effective 5p-start probabilities, and define the cluster total

$$N_{5\mathrm{p}}(C) = \sum_{x \in C} n_{5\mathrm{p}}(x) = \sum_{x \in C} \sum_i \mathrm{Bin}(n_i, p_i(x)).$$

It follows similar to the above that

$$\frac{1}{\mathrm{SNR}} = \frac{\sqrt{\mathrm{Var}[N_{5\mathrm{p}}(C)]_{\mathrm{sampling}}}}{\mathbb{E}[N_{5\mathrm{p}}(C)]} = \frac{1}{\sqrt{\mathbb{E}[N_{5\mathrm{p}}(x)]}}.$$

Summing across a cluster therefore reduces sampling noise by a factor that grows with cluster size. For instance, if a cluster is made of 30 positions each with average count 5, then noise-to-signal ratio for an individual position would be $1/\sqrt{(5)} \approx 0.45$, whereas for the cluster total it would be $1/\sqrt{(30 \cdot 5)} \approx 0.08$.

##### 3. Summed cluster counts provide linear readouts of state abundances

We begin with the model for the observed 5p-start counts in a single cell,

$$n_{5\mathrm{p}}(x) = \sum_i \mathrm{Bin}(n_i,\, p_i(x))\,,$$

where $n_i$ is the abundance of ribosomes in state $i$ and $p_i(x)$ is the effective start probability at position $x$ for state $i$.

Let $C$ be a cluster of positions that share a common relative cutting profile across states. That is, for some reference position $x_0 \in C$,

$$p_i(x) \;=\; \alpha_x\, p_i(x_0) \qquad \text{for all } x \in C \text{ and all states } i,$$

with constants $\alpha_x$ independent of $i$.

The observed cluster total is

$$N_{5\mathrm{p}}(C) = \sum_{x\in C} n_{5\mathrm{p}}(x) = \sum_{x\in C}\sum_i \mathrm{Bin}(n_i,\, \alpha_x p_i(x_0))\,.$$

Each binomial term may be decomposed into its mean plus a fluctuation:

$$\mathrm{Bin}(n_i, q) = n_i q + \varepsilon_i(q), \qquad \varepsilon_i(q) := \mathrm{Bin}(n_i, q) - n_i q.$$

Substituting the decomposition into the cluster total gives

$$N_{5\mathrm{p}}(C) = \sum_{x\in C}\sum_i n_i \alpha_x p_i(x_0) \;+\; \underbrace{\sum_{x\in C}\sum_i \varepsilon_i(\alpha_x p_i(x_0))}_{\text{sampling noise}}\,.$$

Rearranging the first term on the right,

$$\sum_{x\in C}\sum_i n_i \alpha_x p_i(x_0) = \sum_i n_i \left(p_i(x_0)\sum_{x\in C}\alpha_x\right) = \sum_i n_i\, w_{i,C},$$

where we define the cluster–state weights

$$w_{i,C} := \left(\sum_{x\in C}\alpha_x\right) p_i(x_0).$$

The sampling noise term becomes negligible when the cluster size is large. This is because the summation over multiple positions increases signal linearly but noise only as the square root of cluster size. When sampling noise is negligible, the observed cluster total becomes

$$N_{5\mathrm{p}}(C) \;\approx\; \sum_i n_i\, w_{i,C},$$

that is, each cluster provides a readout of a single fixed linear combination of the ribosomal state abundances in each cell. Different clusters will then give readouts of different linear combinations of states. These linear readouts form the inputs to the NMF-based deconvolution performed in the next section.

#### F. Inferring relative ribosomal state abundances via non-negative matrix factorization

Having grouped positions into clusters $C$ that share common relative cutting profiles, and having shown that the summed 5p-start counts in a cluster provide a readout of a particular linear combination of the underlying ribosomal state abundances, we now show how to decompose these linear combinations using non-negative matrix factorization (NMF).

Let $N_{5\mathrm{p}}(C)$ denote the summed 5p-start count for cluster $C$ in a single cell. From the previous section,

$$N_{5\mathrm{p}}(C) \approx \sum_{i=1}^{N_{\mathrm{states}}} w_{i,C}\, n_i,$$

where $w_{i,C}$ are the weights determined by the relative cutting probabilities within the cluster, and $n_i$ are the abundances of ribosomes in state $i$ in that cell.

Collecting all cluster totals for a single cell into a row vector,

$$\mathbf{N}_{5\text{p}}^{\text{cell}} = \big(N_{5\text{p}}(C_1), N_{5\text{p}}(C_2), \ldots, N_{5\text{p}}(C_M)\big),$$

and stacking these vectors across all $K$ cells yields the cell-by-cluster count matrix

$$X \in \mathbb{R}_{\geq 0}^{K \times M},$$

with entries

$$X_{kc} = N_{5\text{p}}^{(k)}(C_c).$$

Under the model above,

$$X \approx NW^{\top},$$

where $N$ contains the ribosomal state abundances for each cell and $W$ contains the cluster-specific cutting weights. Prior to factorization, we normalize $X$ to reduce technical variation. First, each cluster is divided by its mean count across cells,

$$\tilde{X}_{kc} = \frac{X_{kc}}{\langle X_{\cdot c} \rangle},$$

where $\langle X_{\cdot c} \rangle$ denotes the mean of column $c$. Next, each cell is normalized by its total signal,

$$\hat{X}_{kc} = \frac{\tilde{X}_{kc}}{\sum_{c'} \tilde{X}_{kc'}}.$$

NMF is then applied to the normalized matrix,

$$\hat{X} \approx NW^{\top},$$

where

$$N \in \mathbb{R}_{\geq 0}^{K \times N_{\text{states}}}, \qquad W \in \mathbb{R}_{\geq 0}^{M \times N_{\text{states}}}.$$

Column-mean normalization ensures that clusters with inherently higher cut counts do not dominate the factorization, while row-sum normalization removes cell-to-cell differences in total sequencing depth or amplification. Together, these steps highlight relative variations in cluster contributions due to ribosomal state mixtures rather than technical artifacts, improving NMF convergence and interpretability of the recovered matrices $N$ and $W$.

After normalization, we perform NMF on the resulting matrix $\hat{X}$:

$$\hat{X} \approx NW^{\top},$$

where $N \in \mathbb{R}_{\geq 0}^{K \times N_{\text{states}}}$ contains the relative abundances of each ribosomal state in each cell, and $W \in \mathbb{R}_{\geq 0}^{M \times N_{\text{states}}}$ contains the weights. Both $N$ and $W$ are constrained to be non-negative, and the factorization minimizes

$$\min_{N \geq 0,\, W \geq 0} \|\hat{X} - NW^{\top}\|_F^2,$$

where $\|\cdot\|_F$ denotes the Frobenius norm.

The NMF decomposition of normalized cluster totals provides:

- $W$, the cutting weight matrix, representing how each ribosomal state contributes to each cluster.
- $N$, the ribosomal state fraction matrix, giving abundances of each state in each cell, normalized to remove sequencing depth effects.

##### G. Identifying ribosomal states using signature cuts

The NMF decomposition

$$X \approx NW^{\top}$$

is inherently scale non-unique: multiplying a column of $N$ by a constant and dividing the corresponding row of $W^{\top}$ by the same constant does not change the factorization. Moreover, the raw weights $w_{j,C_i}$ returned by NMF have unknown scale that depends on factors such as cluster size and sequence and length biases. Consequently, additional rescaling is required to produce interpretable, fraction-like quantities.

We perform the following steps to obtain a rescaled, interpretable weights matrix:

1. **Un-normalize** $N$: We multiply each row of $N$ by the factor used to normalize the corresponding row of $X$. Because the NMF factors remain scale non-unique, the resulting entries should be interpreted as state activities proportional to the underlying state abundances, up to an unknown state-specific scaling factor.

2. **Compute average inferred state activity**: Let

$$\langle N_j \rangle = \frac{1}{K} \sum_{k=1}^{K} N_{k,j}$$

denote the mean inferred activity of state $j$ across cells.

3. **Rescale** $W$: We multiply each row of $W$ by the corresponding $\langle N_j \rangle$:

$$w_{j,C_i}^{\text{rescaled}} = w_{j,C_i} \langle N_j \rangle.$$

In the underlying model,

$$\mathbb{E}[N_{5\text{p}}(C_i)] \approx \sum_j \langle n_j \rangle \, w_{j,C_i}^{\text{true}},$$

where $\langle n_j \rangle$ denotes the true mean abundance of state $j$. Since the inferred activities $N_{k,j}$ are proportional to the underlying state abundances up to an unknown state-specific scaling factor, $w_{j,C_i}^{\text{rescaled}}$ is proportional to the average contribution of state $j$ to cluster $C_i$ across the cell population, while still reflecting factors such as cluster size and sequence-specific biases.

4. **Column normalization**: Next, we normalize each column of the rescaled weights matrix:

$$\tilde{w}_{j,C_i} = \frac{w_{j,C_i}^{\text{rescaled}}}{\sum_{j'} w_{j',C_i}^{\text{rescaled}}}.$$

As discussed above, the inferred state activities satisfy

$$\langle N_j \rangle = a_j \langle n_j \rangle,$$

where $a_j$ is an unknown positive state-specific scaling factor arising from the non-uniqueness of the NMF decomposition. Consequently,

$$w_{j,C_i}^{\text{rescaled}} \propto \langle n_j \rangle \, w_{j,C_i}^{\text{true}}.$$

The proportionality constants are removed by column normalization, yielding

$$\tilde{w}_{j,C_i} \approx \frac{\langle n_j \rangle \, w_{j,C_i}^{\text{true}}}{\sum_{j'} \langle n_{j'} \rangle \, w_{j',C_i}^{\text{true}}}.$$

Thus, $\tilde{w}_{j,C_i}$ represents the fraction of expected reads in cluster $C_i$ attributable to state $j$, independent of the arbitrary scaling of the NMF factors. This is the blue matrix shown in Fig. (3) of the main text.

4. **Re-normalizing** $N$: To remove the state-specific scaling ambiguity of the NMF decomposition, we first divide each inferred state activity by its average across cells:

$$N_{k,j}^{\text{fc}} = \frac{N_{k,j}}{\langle N_j \rangle}.$$

If $N_{k,j} = a_j n_{k,j}$, where $a_j$ is an unknown state-specific scaling factor, then

$$N^{\mathrm{fc}}_{k,j} = \frac{a_j n_{k,j}}{a_j \langle n_j \rangle} = \frac{n_{k,j}}{\langle n_j \rangle}.$$

Thus, $N^{\mathrm{fc}}_{k,j}$ estimates the fold-change of state $j$ in cell $k$ relative to its population-average abundance.

We then normalize each row:

$$\tilde{N}_{k,j} = \frac{N^{\mathrm{fc}}_{k,j}}{\sum_{j'} N^{\mathrm{fc}}_{k,j'}} = \frac{n_{k,j}/\langle n_j \rangle}{\sum_{j'} n_{k,j'}/\langle n_{j'} \rangle}.$$

The resulting matrix $\tilde{N}$ describes the relative composition of mean-normalized state abundances within each cell. Thus, $\tilde{N}_{k,j}$ should be interpreted as the fraction of the cell's state-abundance fold-change signal attributable to state $j$, rather than as the absolute fraction of ribosomes occupying state $j$.

5. **Signature cuts**: Using the column-normalized weights $\tilde{w}_{j,C_i}$, we define *signature cuts* for state $j$ as clusters satisfying

$$\tilde{w}_{j,C_i} \geq 2 \times \max_{j' \neq j} \tilde{w}_{j',C_i}.$$

These clusters receive a disproportionately large contribution from a single ribosomal state and therefore provide a structural fingerprint for state assignment.

##### H. Cross-experiment comparison of ribosomal state composition

The normalization procedures described above remove the scale ambiguity inherent to NMF and produce quantities that can be compared across experiments.

The normalized weights matrix $\tilde{W}$ describes the fraction of expected reads within each cluster attributable to each ribosomal state. Because MNase concentration directly affects the underlying cutting probabilities, $\tilde{W}$ is expected to vary across digestion conditions. Differences in $\tilde{W}$ therefore reflect changes in cleavage accessibility rather than changes in the underlying abundances of ribosomal states.

In contrast, the normalized state matrix $\tilde{N}$ depends only on the relative abundances of ribosomal states:

$$\tilde{N}_{k,j} = \frac{n_{k,j}/\langle n_j \rangle}{\sum_{j'} n_{k,j'}/\langle n_{j'} \rangle}.$$

Consequently, $\tilde{N}$ is independent of the MNase concentration used to generate the data and should be preserved across experiments performed on the same underlying cell population.

This distinction enables joint analysis of datasets collected at different MNase concentrations. While the cutting-weight matrices $\tilde{W}$ may differ substantially between experiments, the inferred state composition matrix $\tilde{N}$ should remain consistent. In this scenario, no additional normalization is required other than what was described above, provided that the cells in each experiment are sampled from the same underlying population distribution.

A second scenario arises when datasets are generated from populations with different cell-type compositions. In this case, the population-average state abundances $\langle n_j \rangle$ used for normalization may differ between datasets, even if the ribosomal state composition within each cell type remains unchanged. As a result, direct comparison of $\tilde{N}$ can be confounded by differences in cell-type abundance rather than differences in ribosomal state composition.

This situation occurs in our mouse datasets, where the relative abundances of cell types vary between samples. To account for this effect, the normalization factors $\langle n_j \rangle$ are computed using balanced subsamples of cells from each cell type, chosen so that every dataset contains the same relative representation of cell types.

This scale-free representation enables integration of measurements across sequencing runs, experimental batches, and MNase conditions, while separating technical variation in cleavage patterns (captured by $\tilde{W}$) from biological variation in ribosomal state composition (captured by $\tilde{N}$).

##### I. Incorporating 3p-end information

The framework developed above is not restricted to 5p-start positions. For each reference position $x$, one may also define the corresponding 3p-end count, $n_{3\mathrm{p}}(x)$, representing the number of detected fragments whose 3p end maps to position $x$. Throughout, we adopt the convention that a 3p end at position $x$ corresponds to a fragment whose final aligned nucleotide is at position $x - 1$.

Although both quantities are derived from the same fragment population, they capture different aspects of the digestion process. A detected 5p start at position $x$ may be associated with fragments of many different lengths and therefore many distinct 3p ends. Likewise, a detected 3p end may arise from fragments with many different start positions. Thus, 5p starts and 3p ends provide complementary information about the underlying cutting pattern.

For example, a cut at position $x$ followed by a second cleavage at $x + 1$ generates an extremely short fragment that cannot be uniquely mapped and is therefore not detected. In this case, the 5p start at $x$ is absent from the observed data. However, the same cleavage at $x$ may also serve as the 3p end of longer detectable fragments generated from upstream start positions. Consequently, 5p-start counts and 3p-end counts can respond differently to the same underlying cleavage event.

To incorporate both sources of information, we concatenate the 5p-start and 3p-end count vectors for each cell:

$$X_{\text{combined}} = \begin{bmatrix} X_{5\text{p}} & X_{3\text{p}} \end{bmatrix},$$

where the columns of $X_{5\text{p}}$ correspond to 5p-start positions and the columns of $X_{3\text{p}}$ correspond to 3p-end positions.

The covariance-invariant clustering procedure is then applied directly to the combined feature set. Clusters may therefore contain 5p positions, 3p positions, or mixtures of both, provided that they exhibit the same co-variability profile across cells. The resulting cluster counts are then analyzed using the NMF framework described above. In this formulation, both fragment-start and fragment-end information contribute simultaneously to ribosomal state inference.

#### II. Computational workflow for state deconvolution

The computational analyses below were performed in Python (version 3.12.7) using standard scientific packages including NumPy, SciPy, pandas, scikit-learn for data handling and statistical computations. The package igraph with the Leidenalg implementation was used for clustering, and nimfa was used for non-negative matrix factorization implementation.

##### A. Computing single-cell cutting vectors

After mapping and pre-processing the FASTQ files, each single cell is represented by four position-indexed vectors defined on the 45S pre-rRNA reference and rRNA5s reference: 1. The number of 5p cuts at each position along 45s reference. 2. The number of 5p cuts along 5s reference. 3. The number of 3p cuts along 45s reference. 4. The number of 3p cuts along 5s reference.

For each cell $i$, we concatenate these 4 vectors in the above order to form a single cutting profile

$$\text{cuts}_i(x), \qquad x = 1, 2, 3, \ldots, L,$$

where

$$L := 2L_{45\text{s}} + 2L_{5\text{s}} + 2.$$

The additional two positions arise from the 3p-end coordinate convention. Specifically, a 3p end at position $x$ is defined such that the final aligned nucleotide of the fragment lies at position $x - 1$. As a result, a fragment ending at the final nucleotide of the 45S or 5S reference contributes to a 3p-end count at positions $L_{45\text{s}} + 1$ or $L_{5\text{s}} + 1$, respectively.

The first $L_{45\text{s}}$ entries correspond to 5p cuts along the 45S reference, followed by $L_{5\text{s}}$ entries corresponding to 5p cuts along the 5S reference. These are followed by $L_{45\text{s}} + 1$ entries corresponding to 3p cuts along the 45S reference and $L_{5\text{s}} + 1$ entries corresponding to 3p cuts along the 5S reference. Thus, each cell is represented by a unified cutting vector of length $2L_{45\text{s}} + 2L_{5\text{s}} + 2$ containing all MNase cutting information.

For each cell $i$, we computed the normalized cutting frequency as

$$f_i(x) = \frac{\text{cuts}_i(x)}{\sum_{x'} \text{cuts}_i(x')}.$$

##### B. Filtering positions for robust covariance estimation

Our analysis aims to cluster positions according to their normalized covariance profiles. However, the normalized covariance matrix $\eta$ is itself estimated from a finite number of single cells. As a result, positions with low cut counts or weak covariance structure can produce unreliable estimates of $\eta_{xy}$, leading to unstable clustering. To ensure robust estimation of variances, covariances, and ultimately the normalized covariance matrix, we applied the following filters to each position.

1. **Cutting signal abundance.**

   Positions with little or no reproducible signal across the cell population were removed. Specifically, we retained only positions satisfying

   $$\text{median}_i\big(\text{cuts}_i(x)\big) > 0.$$

   Here, $\text{median}_i(\text{cuts}_i(x))$ denotes the median number of cuts observed at position $x$ across all cells. This criterion requires a position to have a nonzero cut count in at least half of the cells and removes positions whose signal is restricted to a small subset of cells.

   For datasets containing multiple cell types, such as the mouse intestine dataset, this filter was applied separately within each annotated cell type. A position was retained if it satisfied the above criterion in at least one cell type. This prevents biologically meaningful cuts specific to a single cell type from being removed when computing the median across the entire dataset.

2. **Finite-cell-count accuracy.**

   Edges in the position graph are constructed only between pairs of positions with similar normalized variances $\eta_{xx}$, as discussed in later Sec. II D. As a result, uncertainty in $\eta_{xx}$ estimates arising from the finite number of cells should be smaller than the $\eta_{xx}$ similarity threshold used during graph construction. Bootstrap resampling was therefore used to estimate the statistical uncertainty of the sampling-noise-corrected diagonal quantity

   $$\eta_{xx} - \frac{\langle f_x / N_{\text{tot}} \rangle}{\langle f_x \rangle^2}.$$

   For each position, 1000 bootstrap datasets were generated by resampling cells with replacement, and the above quantity was recomputed for each bootstrap sample. The standard deviation across bootstrap replicates was taken as the corresponding standard error. Positions were retained only if the relative bootstrap error was below a dataset-specific threshold (20% for the human cell-cycle datasets and 25% for the mouse intestinal datasets), ensuring that sampling uncertainty was smaller than the $\eta_{xx}$ similarity criterion used during graph construction.

3. **Variability bounds.** Positions were required to have normalized variances within dataset-specific ranges ($0.01 < \eta_{xx} < 5$ for the human cell-cycle datasets and $0.01 < \eta_{xx} < 10$ for the mouse intestinal datasets). The lower cutoff removes positions with negligible variability across cells, whereas the upper cutoff removes positions exhibiting unusually large variability that may reflect technical artefacts. The larger upper bound used for the mouse intestinal datasets accommodates the broader range of biological variability observed across the more heterogeneous cell populations.

4. **Covariance-profile quality.**

   Following computation of the normalized covariance matrix $\eta$, positions were further filtered according to the strength and reproducibility of their covariance profiles. First, the 15% of positions with the lowest row-wise sums of squared covariance values,

   $$\sum_y \eta_{xy}^2,$$

   were discarded to remove positions with weak covariance structure. The robustness of this statistic was then assessed by bootstrap resampling of cells. For each bootstrap replicate, the full $\eta$ matrix was recomputed and the row-wise sums recalculated. A signal-to-noise ratio (SNR) was defined as the full-data row-wise sum divided by its bootstrap standard deviation. Only positions with SNR $> 3$ were retained for downstream clustering.

Positions passing all filters formed the set of retained positions $P$, which were retained for $\eta$ estimation and downstream analysis.

##### C. Computing the normalized covariance matrix $\eta$

On the set of retained positions $P$, the normalized cutting frequencies $f_i(x)$ were recomputed per cell. We then computed

$$\langle f(x)f(y)\rangle = \frac{1}{N_{\text{cells}}}\sum_{i}^{N_{\text{cells}}} f_i(x)f_i(y), \qquad \langle f(x)\rangle = \frac{1}{N_{\text{cells}}}\sum_{i}^{N_{\text{cells}}} f_i(x),$$

and the normalized covariance

$$\eta_{xy} = \frac{\langle f(x)f(y) - \langle f(x)\rangle\langle f(y)\rangle}{\langle f(x)\rangle\langle f(y)\rangle}.$$

As discussed in Theory Sec. I D, the diagonal elements $\eta_{xx}$ contain a contribution from sequencing sampling noise. Specifically, Theorem 1 shows that when the relative effective cutting probabilities between two positions $x$ and $y$ are equal, then $\eta_{xz} = \eta_{yz}$ for all $z \neq x, y$, and

$$\eta_{xy} = \eta_{xx} - \frac{\langle f(x)/N_{\text{tot}}\rangle}{\langle f(x)\rangle^2} = \eta_{yy} - \frac{\langle f(y)/N_{\text{tot}}\rangle}{\langle f(y)\rangle^2},$$

where $N_{\text{tot}} = \sum_{x'} \text{cuts}_i(x')$ is the total per-cell cut count. This corresponds to a diagonal correction term $\frac{\langle f(x)/N_{\text{tot}}\rangle}{\langle f(x)\rangle^2}$, which was subtracted from $\eta_{xx}$ for every $x$. The $\eta_{xz} = \eta_{yz}$ relation then holds for all $z$ after subtracting the sequencing-sampling noise term from the diagonal of $\eta$. Finally, the matrix was symmetrized, $\eta \leftarrow \frac{1}{2}(\eta + \eta^\top)$, to remove tiny numerical asymmetries arising from computational floating point errors.

##### D. Clustering the $\eta$ matrix

Leiden clustering was applied to cluster rRNA positions via the normalized covariance matrix $\eta$. Each position $x$ was represented by its row vector

$$\eta_x = (\eta_{x1}, \eta_{x2}, \ldots, \eta_{xL}),$$

which captures the normalized covariance of position $x$ with all other positions. Graph connectivity was determined using the Pearson correlation between covariance profiles together with the normalized covariance criteria below. For position pairs that satisfied these criteria, the Euclidean distance between $\eta_x$ and $\eta_y$ was converted to a similarity score in $[0, 1]$, which was used as the edge weight. An edge between positions $(x, y)$ was drawn only if all of the following conditions held:

$$\rho_{\eta_x,\eta_y} \geq \rho_{\min} \quad \& \quad \left|\frac{\eta_{xx}-\eta_{yy}}{\eta_{xx}+\eta_{yy}}\right| < \delta_\eta.$$

These criteria ensure that positions are connected only if they exhibit similar covariance profiles, both in magnitude and shape. For all datasets, we set $\rho_{\min} = 0.8$. For the RPE1-FUCCI datasets, we used $\delta_\eta = 0.2$ for all MNase concentrations shown in Extended Data Fig. 2 and $\delta_\eta = 0.25$ for the lowest MNase concentration shown in Extended Data Fig. 3, reflecting the reduced number of positions passing the filtering criteria at this digestion level. For the mouse datasets, we used $\delta_\eta = 0.25$. Leiden clustering was then applied to the resulting weighted graph using a fixed random seed and dataset-specific resolution parameters. Clusters containing 20 or fewer positions were discarded.

Leiden clustering was then applied to the resulting weighted graph using a fixed random seed. Because the structure of the $\eta$ matrix depends on the MNase digestion level and the distribution of cell types within a dataset, the resolution parameter was selected separately for each analysis. For the RPE1-FUCCI datasets, a resolution of 12 was used for all MNase concentrations shown in Extended Data Fig. 2, except for the highest MNase concentration, where a resolution of 8 was used to avoid over-partitioning. For the low-MNase dataset shown in Extended Data Fig. 3, a resolution of 5 was used owing to the reduced number of retained positions. For the mouse intestinal datasets, resolutions of 12 and 13 were used for batches 1 and 2, respectively. Clusters containing 20 or fewer positions were discarded.

To assess cluster robustness, we identified cells in which a cluster contributed an unusually large fraction of the total cut signal. For each cluster, the top 1% of cells ranked by cluster signal fraction were removed and the normalized covariance matrix $\eta$ was recomputed using the remaining cells. For datasets containing multiple cell types, this trimming was performed independently within each annotated cell type to avoid preferentially removing cells from abundant cell types that naturally contribute strongly to a given cluster. Let $\eta_{\text{orig}}$ and $\eta_{\text{trim}}$ denote the cluster submatrices before and after trimming, respectively.

$$R_\eta = \frac{|\langle \eta_{\text{orig}} \rangle - \langle \eta_{\text{trim}} \rangle|}{\langle \eta_{\text{orig}} \rangle},$$

where $\langle \eta \rangle$ denotes the mean value of the cluster covariance matrix. Clusters were retained only if $R_\eta < R_{\eta,\max}$, where $R_{\eta,\max} = 0.3$ for the RPE1-FUCCI datasets and 0.15 for the mouse intestinal datasets. The more stringent threshold for the mouse datasets reflects their larger number of cells (∼1000 versus ∼200 per analysis), enabling smaller changes in $\eta$ to be distinguished from sampling noise. This procedure removes clusters whose covariance structure is dominated by a small number of outlier cells.

##### E. Non-negative matrix factorization of summed cluster counts

Clusters retained from the previous step were used to construct the input matrix. For each cell, cuts from positions within the same cluster were summed, giving a matrix

$$X \in \mathbb{R}_{\geq 0}^{K \times M},$$

where $K$ is the number of cells and $M$ is the number of retained clusters. The entry $X_{kc}$ denotes the total number of cuts assigned to cluster $c$ in cell $k$.

###### 1. *Scaling input matrix*

To prevent clusters with large cut counts from dominating the NMF objective, each cluster was first normalized by its average value across all cells:

$$\tilde{X}_{kc} = \frac{X_{kc}}{\langle X_{kc} \rangle}.$$

This puts all clusters on a comparable scale while preserving cell-to-cell variation within each cluster. Next, to account for differences in sequencing depth and overall signal between cells, each row of $\tilde{X}$ was normalized to sum to one:

$$\hat{X}_{kc} = \frac{\tilde{X}_{kc}}{\sum_{c'} \tilde{X}_{kc'}}.$$

The resulting matrix $\hat{X}$ was used as input for NMF.

###### 2. *Non-negative matrix factorization procedure*

NMF was applied to the normalized cluster–by–cell matrix $\hat{X}_{kc}$ using the `nimfa` Python package. The factorization rank $N_{\text{states}}$ was chosen based on the biological complexity of the dataset and the interpretability of the resulting factors. For the RPE1 analysis shown in Fig. 3, we set $N_{\text{states}} = 3$, corresponding to the minimum number of ribosomal states expected a priori: an open state, a closed state, and a preribosomal biogenesis state. For the mouse intestine data, $N_{\text{states}} = 3$ did not recover a distinct state characterized by cuts on the ribosomal outer surface. Increasing the factorization rank to $N_{\text{states}} = 5$ resolved this additional state while preserving the previously identified states, and was therefore used for the intestine analysis. To improve robustness and reduce sensitivity to local minima, we performed a multi-initialization NMF procedure using the `nndsvd`, `random_c`, and `random_vcol` initialization methods provided by `nimfa`. The deterministic `nndsvd` initialization was run once, while each randomized initialization was run 100 times, yielding 201 total NMF fits for each factorization rank. The factorization with the lowest residual sum of squares was retained. All factorizations used the Brunet multiplicative-update algorithm (`method = brunet`) with Kullback–Leibler divergence updates (`update = divergence`) and a maximum of 5000 iterations.

#### III. Mega-SEC Analysis

To quantify the fraction of free large subunits, we combined size exclusion chromatography (Mega-SEC) absorbance measurements with mass spectrometry-derived stoichiometric information. Because the chromatographic peak corresponding to large subunits (60S) contains a mixture of free large subunits and intact monosomes (80S), a de-mixing analysis was required.

##### A. Quantification of total ribosomes

Absorbance measurements at 260 nm and 280 nm were used to estimate RNA concentration while correcting for protein contributions using the Beer–Lambert law [2]. The Beer–Lambert law relates absorbance $A(x)$ at elution position $x$ to concentration:

$$A(x) = \epsilon\, c(x)\, L,$$

where $\epsilon$ is the mass extinction coefficient, $c(x)$ is the concentration of the absorbing species, and $L$ is the optical path length. For a mixture of RNA and protein, absorbance is additive [2]:

$$A_\lambda(x) = \epsilon_\lambda^{\mathrm{RNA}} c_{\mathrm{RNA}}(x) L + \epsilon_\lambda^{\mathrm{P}} c_P(x) L,$$

where $A_\lambda(x)$ is absorbance at wavelength $\lambda$, $c_{\mathrm{RNA}}(x)$ and $c_P(x)$ are RNA and protein concentrations, and $\epsilon_\lambda^{\mathrm{RNA}}$ and $\epsilon_\lambda^{\mathrm{P}}$ are their respective extinction coefficients.

Using measurements at 260 nm and 280 nm, together with representative extinction coefficients for RNA and protein, we solved the system of equations for RNA concentration. Specifically, we used $\epsilon_{260}^{\mathrm{RNA}} = 0.025$ $(\mu\mathrm{g/mL})^{-1}\mathrm{cm}^{-1}$ and $\epsilon_{280}^{\mathrm{RNA}} = 0.0125$ $(\mu\mathrm{g/mL})^{-1}\mathrm{cm}^{-1}$ for RNA, corresponding to standard spectrophotometric conventions that 1 $\mathrm{A}_{260}$ unit corresponds to 40 $\mu$g/mL single-stranded RNA and that pure RNA exhibits $A_{260}/A_{280} \approx 2$. For protein we used $\epsilon_{260}^{\mathrm{P}} = 0.00057$ $(\mu\mathrm{g/mL})^{-1}\mathrm{cm}^{-1}$ and $\epsilon_{280}^{\mathrm{P}} = 0.001$ $(\mu\mathrm{g/mL})^{-1}\mathrm{cm}^{-1}$ [3].

$$c_{\mathrm{RNA}}(x) = \frac{\epsilon_{280}^{\mathrm{P}} A_{260}(x) - \epsilon_{260}^{\mathrm{P}} A_{280}(x)}{L\left(\epsilon_{260}^{\mathrm{RNA}}\epsilon_{280}^{\mathrm{P}} - \epsilon_{280}^{\mathrm{RNA}}\epsilon_{260}^{\mathrm{P}}\right)}.$$

Integrating $c_{\mathrm{RNA}}(x)$ over chromatographic regions corresponding to polysomes, monosomes, and free 60S subunits (via summed areas of exponentially-modified Gaussian fits of raw absorbance profiles) gives the RNA nucleotide abundance inside these fractions. To convert this RNA measurement into ribosome abundance, we express all quantities in equivalent 80S ribosome units:

$$T = p + m + \alpha f. \tag{8}$$

Here, $T$ is the total integrated $c_{\mathrm{RNA}}$ signal, $p$ is the number of ribosomes in polysomes (i.e., total ribosomes bound to mRNA), $m$ is the number of monosomes (80S ribosomes), $f$ is the number of free 60S subunits, and $\alpha$ is a scaling factor accounting for the reduced RNA content and thus absorptivity of a 60S subunit relative to an 80S ribosome.

##### B. De-mixing of the 60S fraction

The 60S chromatographic fraction contains free 60S subunits with some monosome spillover. The integrated absorbance over this region is

$$B = \beta m + \alpha f, \tag{9}$$

where $B$ is the integrated signal over the 60S fraction and $\beta$ is the fraction of total monosomes contributing to this region.

To quantify this mixture, we used mass spectrometry to measure the ratio of summed riBAQ large-subunit (LSU) to small-subunit (SSU) ribosomal proteins (after normalizing by total protein count in each subunit). Because 80S ribosomes contain both LSU and SSU proteins, this ratio is expected to be approximately 1 for pure monosomes, while free 60S subunits contribute only LSU proteins and increase this ratio (see Fig. 4d of main text). The measured ratio is given by $C$:

$$C = \frac{\beta m + f}{\beta m}. \tag{10}$$

Combining Eq. (9) and Eq. (10) yields

$$f = \frac{B(C-1)}{1+\alpha(C-1)}. \tag{11}$$

##### C. Fraction of free 60S subunits

The fraction of ribosomes present as free 60S subunits is defined as

$$F = \frac{f}{p+m+f}.$$

Using Eq. (8) and Eq. (11), we obtain

$$F = \frac{B(C-1)}{T\,[1+\alpha(C-1)] + (1-\alpha)B(C-1)}.$$

Thus, the fraction of free 60S subunits can be estimated from the measurable quantities defined in Eq. (8), Eq. (9), and Eq. (10), together with the parameter $\alpha$. The parameter $\alpha$ was estimated as 0.741 from the relative RNA content of ribosomal subunits, taken as the fraction of total ribosomal RNA contributed by the large subunit, computed as the ratio of the combined nucleotide length of 28S, 5.8S, and 5S rRNAs to the total nucleotide length of all ribosomal rRNAs (28S, 5.8S, 5S, and 18S). We computed $F$ this way independently using mass spectrometry-derived ratios from Fraction 3 and Fraction 4, which both lie within the 60S chromatographic peak.

Note that this analysis involves assumptions and sources of uncertainty. Larger residuals were observed in the 80S peak fits owing to the broader shape. In addition, RNA extinction coefficients were based on standard values for single-stranded RNA. However, folded rRNA may exhibit somewhat different extinction properties, which were not explicitly modeled in this analysis.


[1] G. M. Cooper and K. W. Adams, *The cell: a molecular approach* (Oxford University Press, 2023).

[2] P. Brescia, Micro-volume purity assessment of nucleic acids using a260/a280 ratio and spectral scanning, BioTek Instruments, Winooski, VT, Tech. Rep. AN060112–12, Rev , 06 (2012).

[3] J. Z. Porterfield and A. Zlotnick, A simple and general method for determining the protein and nucleic acid content of viruses by uv absorbance, Virology **407**, 281 (2010).
